# Deep learning-driven characterization of single cell tuning in primate visual area V4 supports topological organization

**DOI:** 10.1101/2023.05.12.540591

**Authors:** Konstantin F. Willeke, Kelli Restivo, Katrin Franke, Arne F. Nix, Santiago A. Cadena, Tori Shinn, Cate Nealley, Gabrielle Rodriguez, Saumil Patel, Alexander S. Ecker, Fabian H. Sinz, Andreas S. Tolias

## Abstract

Deciphering the brain’s structure-function relationship is key to understanding the neuronal mechanisms underlying perception and cognition. The cortical column, a vertical organization of neurons with similar functions, is a classic example of primate neocortex structure-function organization. While columns have been identified in primary sensory areas using parametric stimuli, their prevalence across higher-level cortex is debated, particularly regarding complex tuning in natural image space. However, a key hurdle in identifying columns is characterizing the complex, nonlinear tuning of neurons to high-dimensional sensory inputs. Building on prior findings of topological organization for features like color and orientation, we investigate functional clustering in macaque visual area V4 in non-parametric natural image space, using large-scale recordings and deep learning–based analysis. We combined linear probe recordings with deep learning methods to systematically characterize the tuning of >1,200 V4 neurons using *in silico* synthesis of most exciting images (MEIs), followed by *in vivo* verification. Single V4 neurons exhibited MEIs containing complex features, including textures and shapes, and even high-level attributes with eye-like appearance. Neurons recorded on the same silicon probe, inserted orthogonal to the cortical surface, often exhibited similarities in their spatial feature selectivity, suggesting a degree of functional organization along the cortical depth. We quantified MEI similarity using human psychophysics and distances in a contrastive learning-derived embedding space. Moreover, the selectivity of the V4 neuronal population showed evidence of clustering into functional groups of shared feature selectivity. These functional groups showed parallels with the feature maps of units in artificial vision systems, suggesting potential shared encoding strategies. These results demonstrate the feasibility and scalability of deep learning–based functional characterization of neuronal selectivity in naturalistic visual contexts, offering a framework for quantitatively mapping cortical organization across multiple levels of the visual hierarchy.

## Introduction

From the intricate layering of neurons with diverse functions in the retina (e.g. Masland, 2001) to the topographic maps of the cerebral cortex (e.g. Penfield & Boldrey, 1937), for decades neuroscientists have been pursing the quest to discover general organizing principles that relate the structure (anatomy) and function (physiology) of the brain. For instance, the concept of functional cortical columns, first described by Mountcastle (1957) in the somatosensory cortex and later in the primary visual cortex (V1) by Hubel & Wiesel (1968), represents one such well-established motif, thought to recur across the primate neocortex (discussed in Horton & Adams, 2005). In this canonical organization, neurons with similar function are vertically organized across cortical layers. Considering that the connections within the cortex are locally dense and span the cortical layers, this configuration enables neurons with similar response functions to synaptically interact, thus facilitating computations to transform information within and across the layers (e.g. Cadwell et al., 2020; Campagnola et al., 2022; Jiang et al., 2015)

Obtaining a comprehensive understanding of the relationship between anatomy and function requires a thorough characterization of neuronal stimulus selectivity or tuning. The selectivity of neurons in monkey and cat V1 for simple visual features, such as orientation, phase, or spatial frequency (Issa et al., 2000; Victor et al., 1994), enables the characterization of the tuning properties of these neurons using well-defined parametric stimuli such as gratings. This has greatly facilitated the identification of general organizing principles of neuronal function in the early visual areas of the cortical hierarchy (e.g. Ohki & Reid, 2014). However, neurons in higher visual areas prefer more complex visual features found in natural scenes, such as shapes, textures, objects, and faces (e.g. Bashivan et al., 2019; Kim et al., 2019; Ponce et al., 2019; Tang et al., 2020; Tsao et al., 2003), which are not easily parameterized. The immense diversity and high dimensionality of the natural image space make it challenging to systematically characterize more complex visual function and link it to an organizing structure.

Although prior studies have described feature selectivity in area V4 (e.g. Bashivan et al., 2019; Gallant et al., 1993; Kim et al., 2019; Pasupathy & Connor, 2002; Tang et al., 2020; Wang et al., 2023), less is known about how this selectivity is organized across cortical depth or whether it exhibits topographic structure akin to columnar motifs observed in early visual cortex. While orientation, curvature and color domains have been reported to exhibit spatial clustering within V4 (Conway & Tsao, 2009; Ghose & Ts’o, 1997; Tang et al., 2020; Tanigawa et al., 2010; Wang et al., 2024; Zhang et al., 2023), other work has suggested a more patchy or irregular distribution of feature selectivity (Kotake et al., 2009; Namima et al., 2025), raising debate over whether V4 exhibits consistent columnar or topographic organization. It is therefore unclear whether higher-order tuning, especially under natural image stimulation, follows similar spatial principles. Addressing this question requires scalable, data-driven approaches capable of characterizing neuronal tuning within the rich statistics of natural images, without relying on strong assumptions about feature space.

Advancements in deep learning promise to overcome these challenges. Specifically, recent deep learning functional models of the brain can accurately predict responses to arbitrary stimuli (Bashivan et al., 2019; Cadena et al., 2019; Walker et al., 2019), enabling essentially unlimited *in silico* experiments that complement rather than replace in vivo recordings, and allow exploration of high-dimensional visual feature spaces that are experimentally inaccessible. This can be used for a comprehensive characterization of neuronal tuning function, such as identifying the neurons’ optimal stimuli (Bashivan et al., 2019; Franke et al., 2022; Höfling et al., 2022; Kindel et al., 2019; Walker et al., 2019), map their invariances (Ding et al., 2023b), characterize contextual modulation (Fu et al., 2023) or characterize how multiple distinct tuning properties or nonlinear contextual effects relate to each other (Ustyuzhaninov et al., 2022). The predictions derived from these *in silico* analyses can then be verified through *in vivo* closed-loop experiments, known as inception loops, which have been successfully applied to single neurons in mice (Ding et al., 2023b; Franke et al., 2022; Fu et al., 2023; Höfling et al., 2022; Walker et al., 2019) and populations of neurons in macaque visual cortex (Bashivan et al., 2019).

In this study, we adapted the inception loop paradigm for macaque electrophysiological single-unit recordings to systematically map stimulus selectivity and analyze the structure-function organization of neurons in visual area V4. We used deep neural networks to build an accurate model of >1,200 recorded V4 neurons, capable of predicting responses to arbitrary images and used it to synthesize the most exciting image (MEI) for individual neurons, which we subsequently verified *in vivo*. We found that neurons recorded on the same silicon probe orthogonal to the cortical surface appeared to have similar spatial features compared to MEIs of neurons recorded in different locations, and verified this impression with human psychophysics and a non-linear embedding space based on image similarity. Furthermore, the MEIs formed isolated clusters in the non-linear embedding space, indicating that V4 neurons separate into distinct functional groups that are selective for specific complex visual features, such as oriented fur patterns, grid-like motifs, curvatures, or even high-level attributes reminiscent of eyes. Interestingly, these functional groups closely resemble the feature maps of early-to mid-level units in deep neural networks trained on image classification (Olah et al., 2020), suggesting that computational principles are shared among biological and artificial visual systems. These findings reveal functional clustering within V4 that is consistent with a topographic organization of feature selectivity and support the notion of a columnar-like organization of natural image selectivity in area V4.

## Results

### Deep neural network approach captures tuning properties of individual monkey V4 neurons

To systematically study the neuronal tuning properties of monkey V4 neurons in the context of natural scenes, we combined large-scale neuronal recordings with deep neural network modeling. To this end, we presented natural images to awake, head-fixed macaque monkeys and monitored the spiking population activity of V4 neurons using acute electrophysiological recordings with 32-channel linear arrays spanning 1,920 µm in depth, covering the majority of the 2 mm cortical depth (Fig. 1a; Denfield et al., 2018). In each recording session, we displayed 9,000–12,075 gray-scale images from the ImageNet database (Deng et al., 2009) organized in a trial structure, where each trial consisted of 15 images, each presented for 120 ms, followed by a gray screen 1,200 ms inter-trial period. During image presentation, the monkey was trained to maintain fixation on a fixation spot offset from the center of the monitor (Fig. 1a). The spot’s exact location was selected prior to each recording session such that the neurons’ population receptive field (RF), determined by using a sparse random dot stimulus, was centered on the monitor. Post-hoc spike sorting of the neuronal activity recorded across 100 sessions from two monkeys isolated the single-unit visual activity of 1,224 individual V4 neurons (Fig. 1c), resulting in a large dataset of well-isolated single-unit activity in monkey V4.

**Fig. 1.**
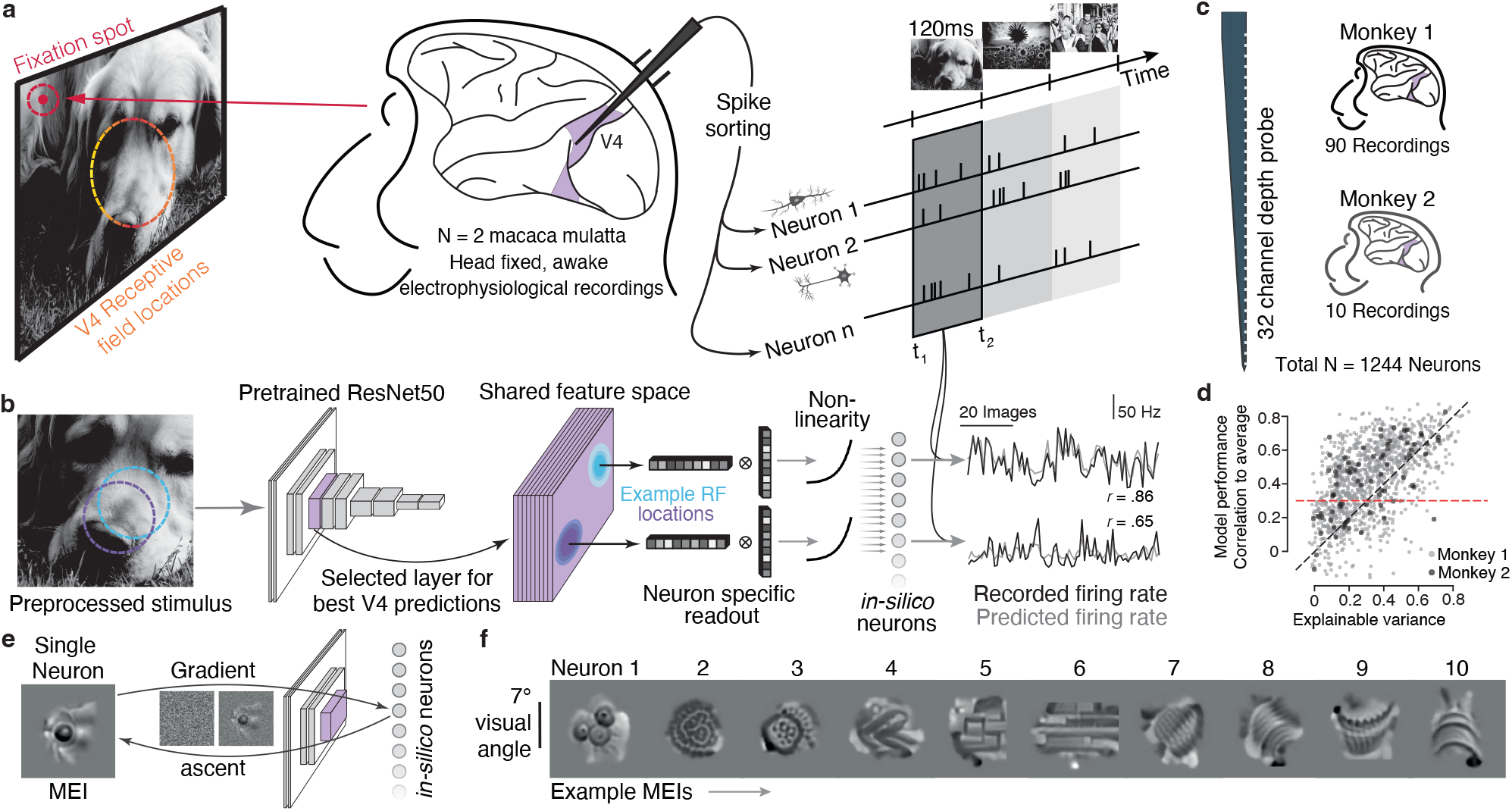
Deep neural network approach captures tuning properties of individual monkey V4 neurons. **a**, Schematic illustrating experimental setup: Awake, head-fixed macaque monkeys were presented with static natural images after fixating for 300 ms (120 ms presentation time per image, 15 images per trial, 1200 ms inter-trial period), while recording the neuronal activity in V4 using 32-channel probes. Animals were fixating on a fixation spot such that the recorded neurons’ population receptive field was centered on the monitor. Post-hoc spike sorting resulted in single-unit activity of individual V4 neurons. **b**, Schematic illustrating model architecture: The pre-processed stimuli (100 x 100 pixels crop) and neuronal responses were used to train a neuron-specific read-out of a ResNet50 pre-trained on an image classification task. Specifically, we selected the ResNet50 layer with the best V4 predictions and computed the neuronal responses by passing the feature activations to a neuron-specific Gaussian readout and a subsequent non-linearity. Traces on the right show average responses (gray) to 75 test images of two example neurons and their corresponding model predictions (black). **c**, Schematic illustrating 32-channels along the probe used for electrophysiological recordings and number of recording sessions per monkey. In total, we recorded the single-unit activity of n=1,244 neurons. **d**, Explainable variance as a measure of response reliability to natural images plotted versus model prediction performance (correlation between prediction and average neural response to repeated presentations) of all cells. Dotted red line indicates a prediction performance of 0.3 used in subsequent analyses (explainable variance mean *±* s.d. = 0.33 *±* 0.19, correlation to average mean *±* s.d. = 0.43 *±* 0.21) **e**, Schematic illustrating optimization of most exciting images (MEIs). For each *in silico* neuron, we optimized its MEI using gradient ascent over n=100 iterations.The whole gray box (full extent) is 14.82°degrees visual angle in width and height. **f**, MEIs of ten example neurons.

To predict the responses of the recorded neurons and characterize the neurons’ tuning properties, we used a deep convolutional neural network (CNN) model (Fig. 1b). Based on previous work in monkey (Bashivan et al., 2019; Cadena et al., 2019, 2022), we used a pretrained goal-directed neural network as a non-linear feature space shared across all neurons and fitted only a simple linear-nonlinear neuron-specific readout (Lurz et al., 2020). Specifically, we chose a robust and high-performing ResNet-50 (Salman et al., 2020) as goal-directed neural network, trained on an image classification task. We selected one of its intermediate layers (layer 3.0) as non-linear feature space because it resulted in the best response predictions of the recorded V4 neurons. This yielded a correlation between response predictions and mean neuronal responses across repetitions of 0.43 (Fig. 1d), indicating that the model reliably captured key tuning properties of the V4 population. In the following, we will refer to this measure as model performance.

Treating our CNN model as a functional digital twin of the population of V4 neurons, we synthesized maximally exciting images (MEIs) (Bashivan et al., 2019; Walker et al., 2019) for individual V4 neurons *in silico* (Fig. 1e). To this end, we optimized a contrast-constrained image to produce the highest activation in the model neuron using regularized gradient ascent. The resulting MEI corresponds to the optimal stimulus of a neuron according to the model, depicting the peak of its tuning curve. We found that MEIs strongly differ across neurons, indicating selectivity for distinct stimulus features like texture, curvature and edges (Fig. 1f), which resemble the features found in the MEIs of V4 multi-unit activity (Bashivan et al., 2019). Our MEIs were also consistent with tuning properties of macaque V4, such as shape, curvature, and texture selectivity, previously, identified using parametric stimuli (e.g. Kim et al., 2019; Pasupathy & Connor, 2001; Pasupathy et al., 2020). In contrast to these previous studies, our data-driven approach enables characterization of single-neuron selectivity in a natural image context, without requiring prior assumptions about feature space or pre-selection of stimuli.

### Closed-loop paradigm verifies model-derived optimal stimuli of single V4 neurons

To demonstrate the model’s accuracy and empirically validate that the synthesized MEIs effectively activated the recorded neurons, we developed a closed-loop paradigm for acute electrophysiological recordings of single neurons (Fig. 2a). Specifically, after fitting the readout for single-unit responses recorded in a “generation” session where natural images were shown, we selected the best six units based on model prediction for *in vivo* verification, generated their MEIs and presented them back to the animal on the same day while recording from the same neurons in a “verification” session. Single units were matched across the generation and verification session using spike waveform similarity and functional consistency of responses to the same natural images (Suppl. Fig. 1). As a control stimulus for each selected unit, we presented the seven most exciting natural image crops identified by the model by screening 5,000 natural images not used during model training. Each crop was matched to the size, position, and contrast of the MEI of a particular neuron. Control stimuli qualitatively resembled the MEIs (Fig. 2b), confirming that modelsynthesized features reflect elements commonly found in natural scenes.

**Fig. 2.**
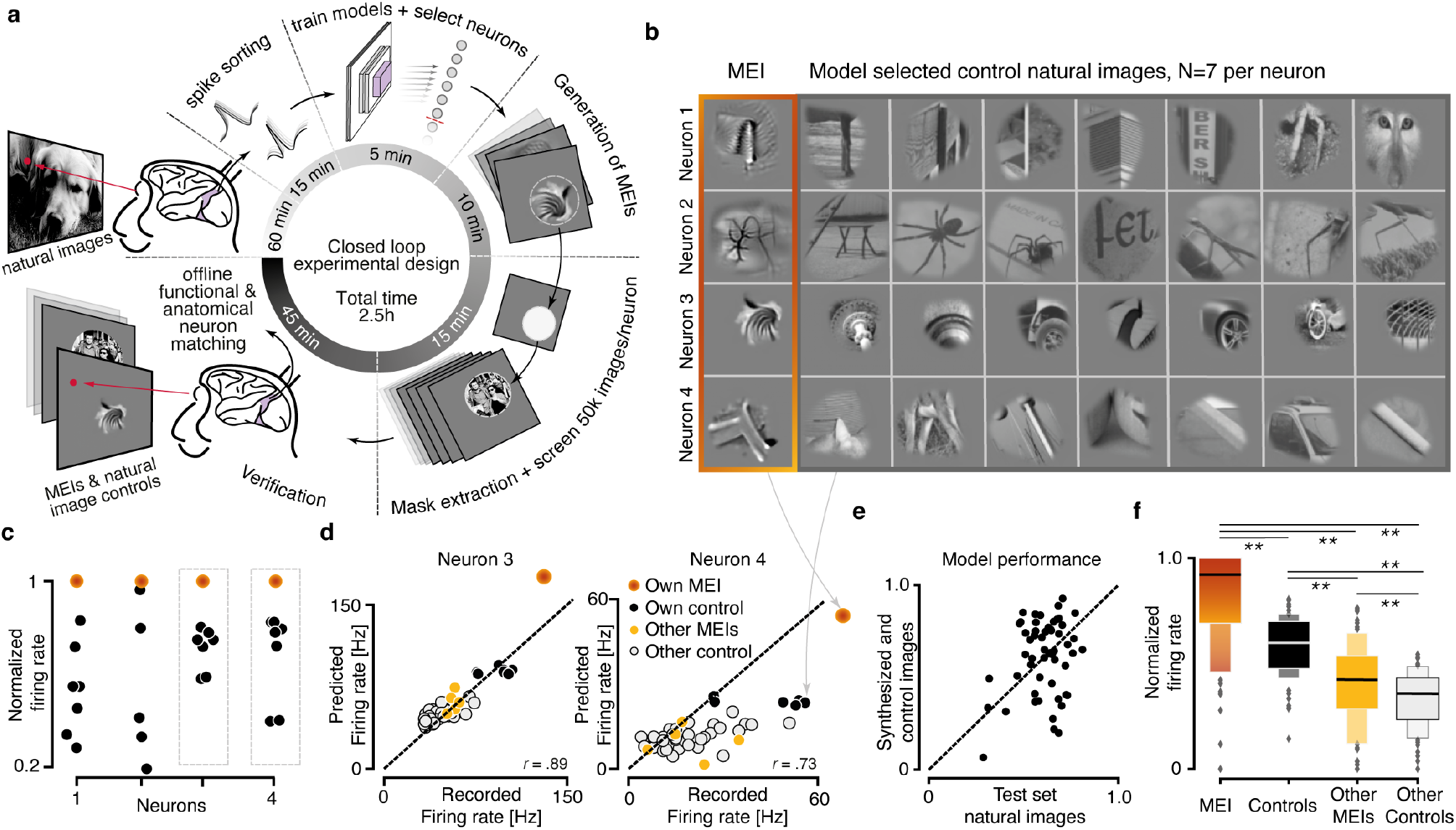
Closed-loop paradigm verifies model-derived optimal stimuli of single V4 neurons. **a**, Schematic illustrating closed-loop experimental paradigm for acute recordings in monkey V4. In brief, after recording and spike sorting of the “generation session”, we train a model, select neurons for experimental confirmation, generate MEIs and identify the most exciting natural image control stimuli, and present both MEIs and controls back to the animal while recording from the same neurons in the “verification session”. Functional and waveform matching of units across recordings is performed offline. **b**, MEI and the seven most exciting natural image crops, selected from 5,000 natural images, for four example neurons. Natural images were matched in size, location and contrast to the MEI. **c**, Peak-normalized recorded responses of the neurons in (b) to their MEI (orange) and control images (black; mean across n=20 repeats). **d**, Recorded versus predicted neuronal activity of two example neurons to their MEI and control stimuli, as well as to MEIs and control stimuli of other neurons. **e**, Scatter plot of model performance on the test set of natural images and the closed-loop stimuli (as shown in d, but for all neurons). Correlation to average: mean *±* s.d. = 0.61 *±* 0.11; Synthesized and selected stimuli: mean *±* s.d. 0.61 *±* 0.20, *n* = 55 neurons. A paired t-test showed no significant difference *p* = .61. **f**, Distribution of peak-normalized mean responses to each neuron’s MEI and control stimuli, as well as MEIs and control stimuli of other neurons for all closed-loop neurons (*n* = 55 neurons, *n* = 24 sessions, *n* = 1 monkey). P-values for a paired t-test are: MEI-Control, 3.22e-08; MEI-OtherMEIs, 2.57e-14; MEI-OtherControls, 3.06e-19; Control-OtherMEIs, 2.86e-07; Control-OtherControls, 1.62e-19; OtherMEIs-OtherControls, 2.99e-05. P-values were corrected for multiple comparisons with Bonferroni correction.

Overall, the model predicted V4 neuronal responses to both full-field natural images and synthesized MEIs with high reliability. Despite the high structural similarity of MEI and control images, the MEI consistently elicited higher neuronal responses than the control images, as well as MEIs and control images of other neurons (Fig. 2c,d,f), indicating neuron-specific selectivity captured by the model. In addition, the model accurately predicted the neurons’ average responses to their own MEIs, to control stimuli, and to the MEIs and control stimuli of other neurons of the same session (two example neurons in Fig. 2d). The absolute scale of the firing rate predictions did not always perfectly match the recorded firing rate, likely due to slow drifts in overall firing rates of some neurons (e. g. Fig. 2d, right). Nevertheless, across neurons, the model trained on the generation session generalized robustly to the verification session, with no significant difference in prediction performance between full-field natural images and synthesized MEIs (*ρ* = 0.61, Fig. 2e). Moreover, the amplitude of neuronal responses to control stimuli and MEIs of other neurons only slightly differed, suggesting that both sets of stimuli share similar low-level visual statistics despite the neuron-specific optimization of MEIs (Fig. 2f).

### Topographic organization of model-derived feature selectivity in macaque V4

Studying how visual selectivity is organized in a particular brain area has revealed key principles of vision, including the pinwheel of orientation columns in primary visual cortex (Bonhoeffer & Grinvald, 1991). In monkey V4, neurons are tuned to more complex visual features like shape and texture (Kim et al., 2019; Pasupathy & Connor, 2001; Pasupathy et al., 2020; Srinath et al., 2020), but it remains unclear whether V4 tuning properties are organized in a columnar manner (Ghose & Ts’o, 1997; Hatanaka et al., 2022; Namima et al., 2025; Tang et al., 2020). We next asked whether our data-driven approach could reveal topographic patterns in feature selectivity, as inferred from model-derived optimal stimuli.

We noticed that MEIs from individual sessions tended to exhibit higher mutual visual similarity compared to MEIs from other sessions (Fig. 3a,b). While the range of preferred stimuli we found spanned a large variety from oriented and comb-like patterns, to grid-like motifs and highlevel attributes, the perceived variability within many sessions was much smaller than across sessions. For example, most neurons in an example session preferred oriented and comb-like patterns (Fig. 3a), while neurons from other example sessions preferred curved edges (session 2 in Fig. 3b) and grid-like patterns (session 3 in Fig. 3b).

**Fig. 3.**
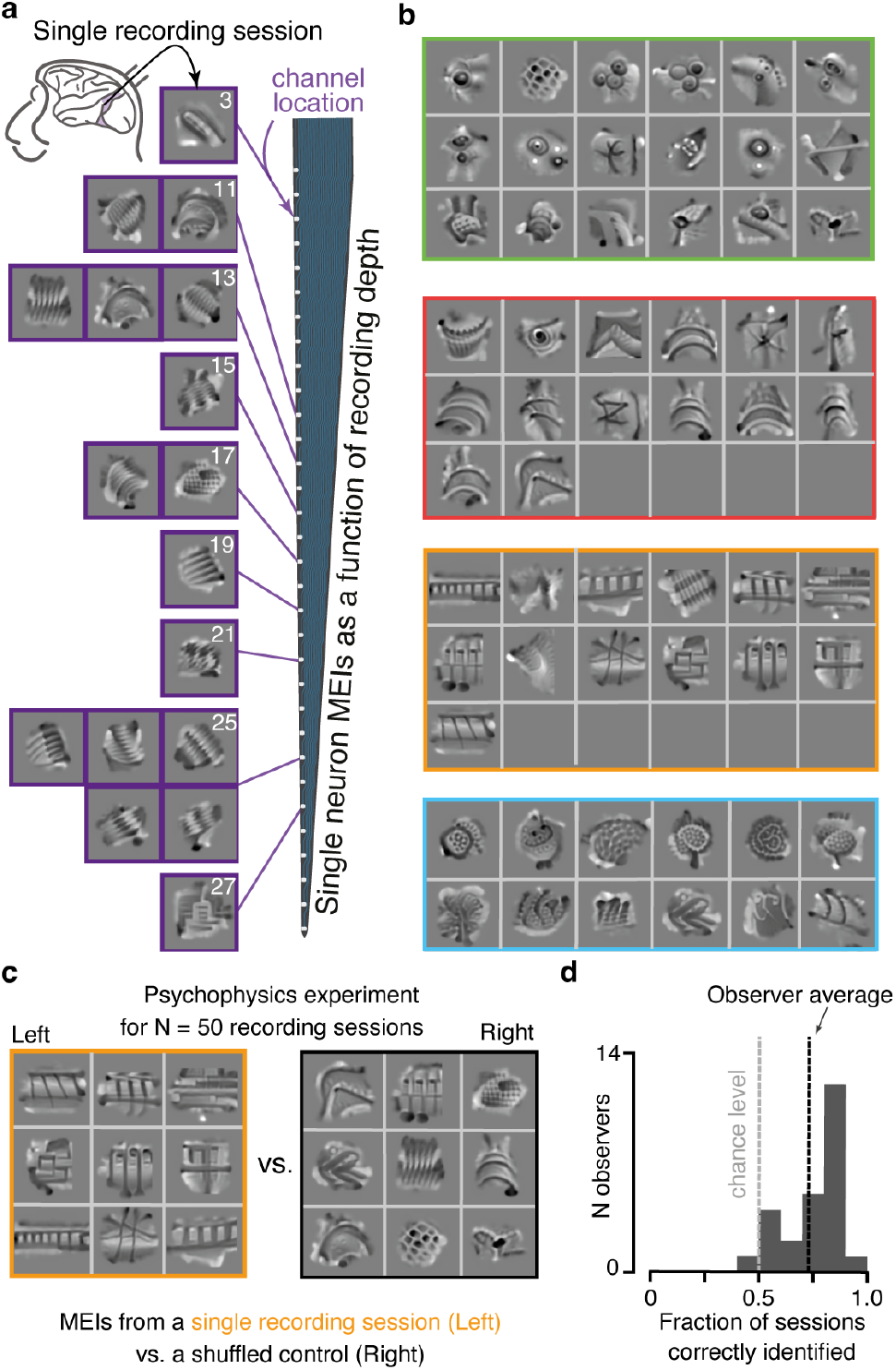
Topographic organization of model-derived feature selectivity in macaque V4. **a**, MEIs of 17 neurons recorded in a single experimental session, arranged according to each neuron’s channel location along the recording probe. Numbers indicate channel, with higher channel numbers meaning greater recording depth. **b**, MEIs of varying numbers of neurons for four different sessions (indicated by different colors). **c**, Schematic illustrating paradigm of simple psychophysics experiment. In one trial, subjects were presented with MEIs of 9 neurons recorded within one session (left) or randomly sampled from all neurons except the target session (right), and reported the location (left or right) of the set of MEIs that looked more consistent (i.e. shared the same image features). The experiment included n=50 trials/sessions. **d**, Distribution of fraction of sessions correctly identified across n=25 observers, with change level and observer average indicated by dotted lines. Mean across subjects, = 0.73; subject-variability in s.d., = 0.13; session-variability in s.d., = 0.21.

To quantify the perceptual similarity of MEIs within a session, we performed a simple psychophysics experiment, where human observers were presented with two sets of MEIs: (1) a selection of nine neurons recorded within one session and (2) a set of nine neurons randomly sampled across sessions. The two sets were presented side-by-side (as shown in Fig. 3c, but without the colored frames), with each set being shown on the left or right at random. The observers then had to report in a two-alternative forced-choice paradigm which set of MEIs looked perceptually more similar. On average, the observers classified MEIs of the same session as being more consistent than a random set of MEIs from different sessions for 73% of the sessions (Fig. 3d), indicating that neurons recorded along the same electrode penetration tend to share similar model-derived feature preferences. Because neurons recorded within a session are arranged roughly along the vertical axis of cortex, these results suggest a local topographic organization of feature selectivity in macaque V4. This pattern is consistent with columnar-like functional clustering but does not constitute direct anatomical evidence for columns.

To further quantify whether tuning properties of V4 neurons are functionally organized in a topographic manner, we performed nonlinear dimensionality reduction to embed the MEIs in a two-dimensional space based on model-derived image feature similarity rather than recording session identity.. In contrast to V1 neurons, whose tuning properties can be compared along clearly defined axes such as orientation or spatial frequency, the complex MEI structure of V4 neurons makes it challenging to quantify the tuning similarity between neurons. To resolve this problem, we used an unsupervised deep learning technique that learns a two-dimensional image-embedding based on mutual similarity of images (Böhm et al., 2023). The model is trained using a contrastive objective: embeddings of augmented versions of the same image are attracted to each other, while embeddings of distinct images are repelled (Fig. 4a). The data augmentations used to generate image variants determine which transformations the model learns to ignore as irrelevant differences. By using random rotations, shifts, and scaling as augmentations, we ensured that MEIs differing only by these geometric transformations would cluster together in the embedding space (Fig. 4a) We chose these augmentations because they generally preserve the identity of many mid- and high-level image features: for instance, an eye remains an eye even after rotating, shifting or scaling. Moreover, we observed that multiple MEIs of the same neuron, generated by starting from different initial noise images during optimization, often exhibited variations in some or all of these dimensions (Suppl. Fig. 2). Since the training of the model is exclusively based on image identity, it does not provide any information about which recording session a particular MEI originated from. Therefore, any grouping or clustering of MEIs by session in the learned embedding space reflects only their visual similarity and not any explicit experimental labels

**Fig. 4.**
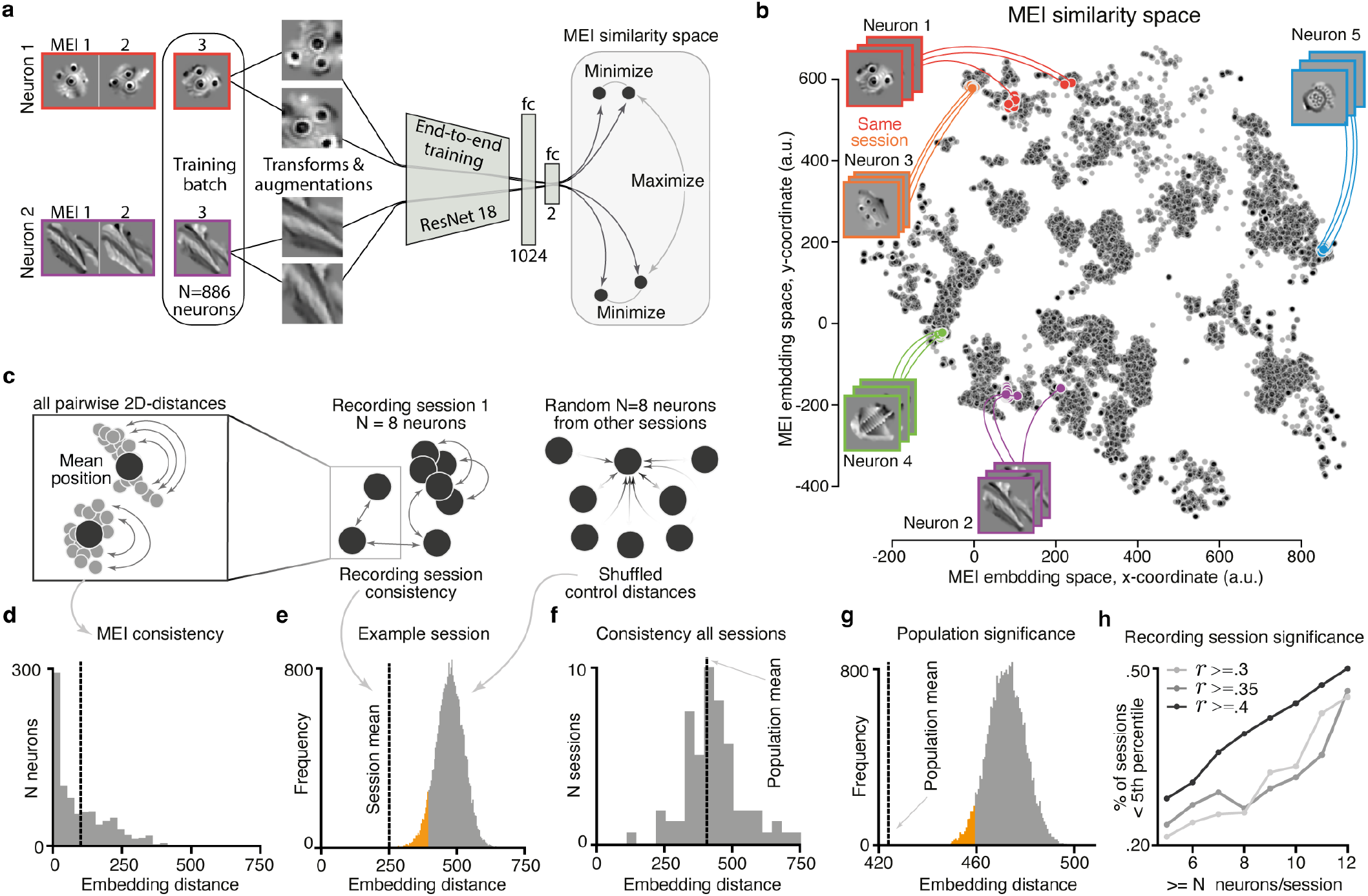
Contrastive clustering of MEIs confirms topographic organization of V4 visual tuning selectivity. **a**, Schematic illustrating contrastive learning approach to quantify MEI similarity. Per neuron, we optimize n=50 MEIs initialized with different random seeds, then select highly activating MEIs, and use one MEI per neuron (n=889) as a training batch. Each MEI is transformed and augmented twice and the model’s objective then is to minimize the distance in a 2D MEI similarity space between different transforms of the same MEI, while maximizing the distance to MEI transforms of other neurons. **b**, Position of all highly activating MEIs (n=19688) of n=889 neurons in a 2D MEI similarity space, with MEIs of five example neurons indicated in different colors. Dots of the same color indicate MEIs optimized from different random seeds of the same neuron. **c**, Schematic illustrating analyses performed on the 2D MEI similarity space. We computed the pairwise 2D distances across all MEIs of one neuron to estimate MEI consistency (left), and all pairwise distances across MEIs of the same recording session to estimate recording session consistency (right). For the latter, we used the distances across a random selection of neurons from other sessions as control. **d**, Distribution of embedding-distances across MEIs of the same neuron. Vertical dotted line indicates mean of the distribution. **e**, Mean distance across neurons from one example session (vertical line), with a null distribution generated by bootstrapping distances across the same number of neurons randomly sampled from all other sessions. Orange shading indicates values <5% percentile. Note that the null distribution depends on how many neurons were recorded in each session, as it estimates the standard error of the mean for each session. **f**, Histogram of session means like in e), but for all sessions. Grand mean across all sessions is indicated by the vertical line. Mean = 423.97 *±* 105.20 s.d. **g**, Mean within-session distance across all sessions from f) along with the mean null distribution across sessions in gray. The population mean significantly deviates from the null distribution (*p <* 4 *×* 10^*−*5^; 25,000 bootstrap samples). Orange shading indicates values <5% percentile. **h**, Percentage of sessions with the within-session distance <5% percentile of the null distribution for different numbers of neurons per session (x-axis) and different model predictions thresholds (shades of gray). The percentiles obtained from the embedding space (including all neurons above a prediction threshold of 0.3) were significantly correlated with the observer agreement (percent correct) of the psychophysics experiment (*ρ* = *−*0.33, p=.019, n=50 sessions).

The resulting embedding space placed neurons with similar MEIs close to each other (neurons 1 and 3 in Fig. 4b) and neurons with different MEI features far away (neurons 1 and 2). Similarly, multiple MEIs of the same neuron, generated by starting from different initial noise images during optimization, were placed nearby in the embedding space as well (groups of the same color in Fig. 4b and Fig. 4c,d). These observations suggest that the model indeed learned to embed MEIs based on image similarity. We next quantified whether neurons recorded within one session share tuning properties as suggested by the observed MEI similarity within sessions (cf. Fig. 3). To this end, we computed the mean pairwise distance in the embedding space across neurons from one session and compared it to a null distribution of distances obtained by computing the mean pairwise distance across randomly picked neurons from different sessions (Fig. 4c,e). While the within-session distance varied across sessions (Fig. 4f), on a population level, it was significantly smaller than the across-session distance (Fig. 4g). The percentage of sessions that showed a significantly smaller distance in MEI similarity than the null distribution increased with higher numbers of neurons recorded per session and with higher prediction performance of the model (Fig. 4h). For more than 12 neurons per session and with the highest performance threshold (correlation to average >0.4), half of the sessions displayed a significantly smaller withinthan across-session distance, indicative for high similarity of MEIs. Importantly, the within-session distances estimated based on the embedding space significantly correlated with the observer agreement (percent of observers who correctly classified a specific session) from the psychophysics results (*ρ* = *−*.33, p=.019), suggesting that MEI distance in the embedding space is informative about MEI perceptual similarity. We additionally confirmed that MEIs of neurons recorded within one session are more similar to each other than to MEIs of neurons recorded in other sessions using an independent similarity metric, namely the representational similarity of MEIs in neuronal response space, which closely mimics perceptual similarity (Kriegeskorte, 2008) (Suppl. Fig. 3). Taken together, these analyses demonstrate consistent functional clustering of tuning properties among V4 neurons recorded along the same cortical penetration. This pattern is consistent with a topographic or columnar-like organization of feature selectivity across cortical depth, as inferred from model-derived representations.

### V4 neurons cluster into distinct functional groups that resemble feature maps of artificial vision systems

At the level of retinal ganglion cells, neurons cluster into specific functional groups or output channels (Baden et al., 2016; Goetz et al., 2022). Whether a similar functional clustering persists for cortical neurons is still an open question. For instance, in V1 it is not clear whether the distinction between simple and complex cells represents two ends of a continuous spectrum or two discrete categories (Mechler & Ringach, 2002). Previous results in mouse primary visual cortex suggest that neurons cluster according to function, but not in an entirely discrete manner (Ustyuzhaninov et al., 2019). Motivated by the structure of the MEI embedding space that exhibited isolated “islands” of MEIs (cf. Fig. 4b), we asked whether tuning properties of V4 neurons fell into similar functional groups, characterized by their preferred stimulus. To address this question, we applied hierarchical clustering (DBSCAN; McInnes et al., 2017) to the two-dimensional MEI embeddings, yielding 17 distinct functional groups (Fig. 5). MEIs within each group exhibited high perceptual similarity and shared stimulus preferences—for instance, neurons in group 11 all responded preferentially to patterns resembling eyes. Conversely, neurons in different groups showed markedly different feature selectivity, particularly for groups positioned distantly in embedding space. For example, while group 11 MEIs contained eye-resembling patterns, group 3 and group 8 MEIs displayed grid-like and comb-like textures, respectively

**Fig. 5.**
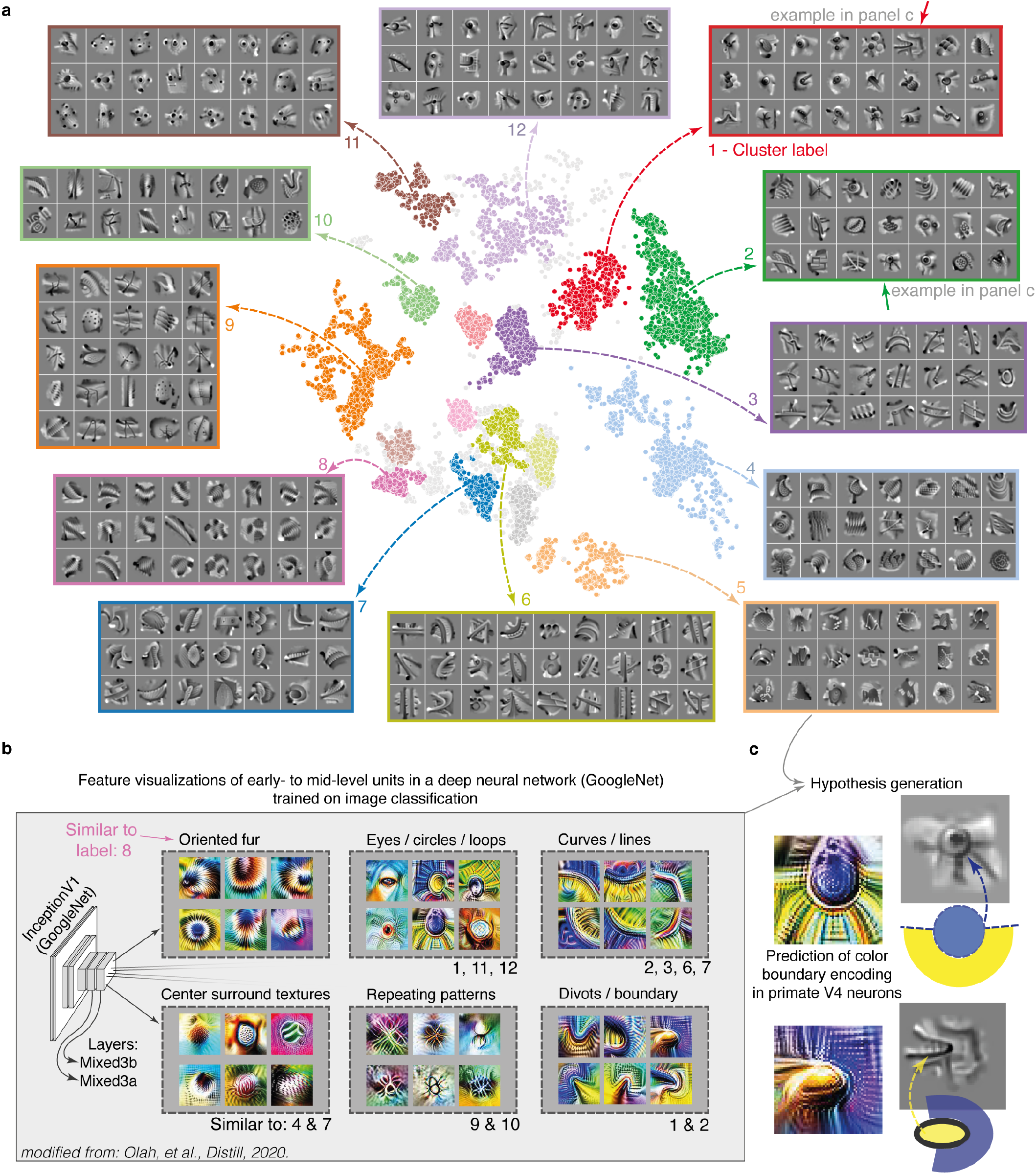
V4 neurons cluster into distinct response modes that resemble feature maps of artificial vision systems. **a**, Position of all highly activating MEIs (n=19,688) of n=889 neurons in the 2D MEI similarity space, color coded based on cluster assignment obtained from the hierarchical clustering algorithm HDBSCAN. For n=12 clusters, we show a random selection of MEIs of different neurons assigned to this cluster. For examples of the other clusters, see Suppl. Fig. 4 and for independent verification of the clusters, see Suppl. Fig. 3. Light gray dots indicate MEIs that could not be assigned to any of the clusters with high probability. **b**, Feature visualizations of early- to mid-level units in the deep neural network InceptionV1 trained in an image classification task (Olah et al., 2020). Units are grouped into distinct categories based on (Olah et al., 2020), with clusters from (a) resembling these categories indicated below. **c**, Example units of the neural network trained on image classification compared with example MEIs exhibiting similar spatial patterns. The resemblance between the two can be used to generate hypotheses, such as to predict color boundary encoding in primate V4 neurons, that can be subsequently tested experimentally.

To ensure that the group structure in the embedding space is not an artifact of the contrastive neighborhood embedding method, we additionally used an independent method to compare within-group to across-group similarities. To that end, we computed the representational similarity of the MEIs using the predicted neuronal responses from our model. Specifically, we centered the receptive fields of all neurons in the model and compared the similarity of two MEIs via the cosine of the predicted population response vectors. Importantly, this similarity metric is unrelated to the embedding space used for clustering and, therefore, provides an independent verification of the identified functional groups. We found that MEIs of neurons assigned to the same group were significantly closer to each other in the neuronal response space than MEIs of neurons assigned to different groups (Suppl. Fig. 3): The within-group similarity in neuronal response space was significantly higher than the across-group similarity for all of the 17 functional groups.

Interestingly, the MEIs of the identified V4 clusters closely resembled the feature visualizations of single units in modern deep neural networks trained on image recognition tasks. For example, a similar preference for specific complex features can be found in the early layers, specifically layer mixed3a, of the InceptionV1 deep network (Olah et al., 2020; Szegedy et al., 2015). This alignment between V4 and deep network features is in line with previous results that found boundary selective units in the AlexNet deep network (Krizhevsky et al., 2012; Pospisil et al., 2018), similar to boundary neurons in monkey V4 (Pasupathy & Connor, 2002). Olah et al. (2020) manually grouped the feature visualizations into different categories like “Oriented fur,” “Eyes/circles/loops” and “Divots/boundaries” (Fig. 5b). These categories exhibit perceptual similarities to the functional groups of V4 neurons identified through hierarchical clustering, suggesting potential convergence in encoding strategies between primate and artificial vision systems. The correspondence between V4 neuronal selectivity and deep neural network features provides a basis for generating hypotheses about visual tuning properties in primate V4 beyond spatial patterns. For example, given that the artificial vision systems were trained on color images, their feature visualizations also characterize color tuning and may inform predictions about color boundary encoding in monkey V4 functional groups (Fig. 5c), amenable to verification through *in vivo* experiments. These comparisons highlight how AI models and neuroscience can inform each other, with artificial representations guiding hypotheses while neural data constrain and refine computational models.

## Discussion

Our work provides evidence for a topographic and columnar-like organization of tuning to spatial patterns in visual area V4, derived from detailed single-neuron tuning properties. This organization was revealed using an iterative image synthesis approach based on a deep neural network model of neuronal activity. This model, serving as a digital twin of V4, enabled characterization of spatial pattern tuning without imposing prior parametric assumptions about feature selectivity. By combining this approach with a learned non-linear image-similarity embedding space and human psychophysics, we found that MEIs from neurons recorded along the same cortical penetration tended to be more similar to each other than those from randomly selected neurons, and that V4 neurons formed functionally coherent clusters in the embedding space. Together, these findings indicate that despite the complexity of V4 stimulus preferences, feature selectivity is locally clustered in a manner consistent with columnar-like or topographic organization.

### Are our MEIs consistent with previous findings?

The idea to use neuronal encoding models to synthesize optimal stimuli for the brain is well established (Lehky et al., 1992) and has already successfully been used in mouse primary visual cortex (Franke et al., 2022; Walker et al., 2019), mouse retina (Höfling et al., 2022), and also macaque V4 (Bashivan et al., 2019). Additionally, there exist deep learning-based methods for synthesizing optimal stimuli which do not rely on neuronal encoding predictive models. Instead, these methods utilize a genetic algorithm to search through the space of a generative deep neural network of images **?**. Many of our MEIs contain features, such as different types of textures, that qualitatively resembled those found by Bashivan et al. (2019). However, some of our MEIs showed shape-like features, including curved strokes, corners, and even higher-level attributes such as individual eye-like stimuli (cluster 7 in Fig. 5a), which have been less emphasized in previous work.

One distinction between our study and that of Bashivan et al. (2019) is that we used silicon probes, enabling us to separate spikes from individual neurons, while they employed chronically implanted Utah arrays and optimized stimuli for single ‘sites’ (multi-units) that likely comprise a mix of multiple neurons. Consequently, it remains unclear from their study whether the complexity of the MEIs stems from the mixing of spikes from multiple neurons or if single neurons already prefer such intricate spatial patterns as displayed by the MEIs. Additionally, MEIs for multi-unit activity could average out specific features like shapes. By isolating and verifying MEIs at single-cell resolution, our results demonstrate that the diversity and complexity of optimal stimuli are already present at the level of individual neurons in macaque V4.

Our MEIs are also consistent with previously described tuning properties of V4 neurons. V4 is part of the ventral pathway which plays a major role in object and shape recognition (Felleman & Van Essen, 1991; Mishkin et al., 1983). Previous studies have shown that V4 cells are selective for complex shapes (Kobatake & Tanaka, 1994), and be tuned to convex and concave shapes of object boundaries at specific locations in the visual field (Gallant et al., 1993; Pasupathy & Connor, 2001). In addition to shape, V4 neurons are also known to be selective to texture (Kobatake & Tanaka, 1994). Kim et al. (2019) found that tuning of single V4 neurons can be placed along a continuum from strong tuning for boundary curvature of shapes to strong tuning for perceptual dimensions of texture. Consistent with these findings, our MEIs captured a wide range of feature preferences. On the one hand, we find MEIs that clearly exhibit curvature elements (e.g. cluster 7 in Fig. 5a) or ‘eye’-like elements (cluster 1). On the other hand, many MEIs have a texture component such as ‘fur’ (cluster 8), dots (cluster 11), or grid-like elements (cluster 6). Interestingly, generating multiple MEIs from different starting points – known as Diverse Exciting Images (DEIs; Cadena et al., 2018; Ding et al., 2023b) – resulted in multiple MEIs that had similar shape and texture features (Suppl. Fig. 2), indicating that single cells in V4 are neither texture- nor shape-invariant. Together, our findings extend previous work by providing a single-cell, data-driven view of selectivity in V4 under naturalistic conditions.

### Topological organization of MEIs in V4

One advantage of our recordings is that we can record simultaneously across cortical layers using silicon probes. This enabled us to characterize the vertical organization of the tuning selectivity to complex spatial patterns in area V4 and to assess whether tuning similarity is consistent with columnar or topographic organization. The presence of columns in V4 has been a matter of debate and controversy, predominantly studied in the domain of color (Kotake et al., 2009) or orientation (Ghose & Ts’o, 1997) because stimuli for these domains are more accessible to low dimensional parametrization. For instance, in the color domain, some studies has reported columnar organization (Conway & Tsao, 2009; Kotake et al., 2009; Tanigawa et al., 2010), while another found no evidence (Tanaka et al., 1986). Beyond that, other studies using natural images or parametric stimuli for curvature found local clustering of similarly tuned neurons (Hatanaka et al., 2022; Tang et al., 2020; Tanigawa et al., 2010; Wang et al., 2024). However, since these studies focused mainly on neurons from superficial layers, it remains unknown whether such topographic clustering extends across cortical depth.

Moreover, it is well-established that V4 neurons exhibit selectivity for complex spatial features (Oliver et al., 2024; Wang et al., 2024) that extend beyond those characterized by parametric stimuli defined by orientation, or parameterized curvature (e.g. Tang et al., 2020). Consequently, in the absence of a detailed identification of the non-parametric optimal stimuli, assessing the hypothesis of functional columns proves to be challenging. For example, if neurons are tuned to similar grid-like textures but with different orientations, using grating stimuli (Ghose & Ts’o, 1997) or predetermined parameterized curvature stimuli (Tang et al., 2020) will obscure the true underlying organization. Our deep learning based image synthesis method avoids these challenges by identifying most exciting stimuli in the high-dimensional pixel space.

To compare MEIs regardless of their spatial complexity, we used human psychophysics and deep learning techniques to assess the similarity between them, specifically employing contrastive learning to create a non-linear embedding based on similarity among MEIs (Böhm et al., 2023). We discovered that neurons across layers recorded in orthogonal penetrations to the cortical surface had MEIs that were perceptually more similar than those from randomly sampled neurons. However, the strength of this clustering effect varied across recording sessions, and was not equally evident in all cases This variability in effect size could have been caused by several factors. First, despite best efforts, our electrode penetrations may not have been perfectly orthogonal to the cortical surface, because the electrodes were aligned relative to the recording chamber. The recording chamber was implanted such that its center was orthogonal to the surface of the cortex. Since the cortex is curved, penetrations further away from the center may not have been perfectly orthogonal to the cortical surface. The brain may also have moved or have been slightly compressed during insertion, potentially resulting in slightly angled penetrations. Second, if there is a topographical organization in V4, it may feature both homogeneous zones and regions with more mixed selectivity, analogous to pinwheels in orientation maps in V1. Thus, we expect a certain fraction of penetrations to be close to such heterogeneous zones. Because extracellular electrodes record the activity of neurons in a roughly cylindrical region around the electrode, we expect a fraction of penetrations to exhibit mixed tuning even if the penetrations were perfectly vertical. Estimating what fraction of penetrations should exhibit consistent tuning is difficult, because the size of the columns and the relationships between them are not yet understood. we estimate that approximately half of the penetrations show a statistically significant bias toward certain feature selectivities, consistent with a topographic organization. While our findings point toward a columnar-like organization of functional selectivity in V4, this interpretation contrasts with recent Neuropixels recordings that revealed sparse, non-columnar clusters for shape and texture tuning (Namima et al., 2025). One possibility is that such discrepancies arise from differences in stimulus complexity, suggesting that columnar organization in V4 may depend on the specific feature space analyzed.

To more accurately delineate and map the topological organization of spatial form tuning in V4, future studies need to combine recording techniques like two-photon functional imaging with deep learning and inception loop approaches. To this end, a parallel study Wang et al. (2024) used wide-field and two-photon calcium imaging across the surface of V4 and identified a topographical map of natural stimulus preference. Their 2D map contained distinct (clustered) functional domains preferring a variety of natural image features, ranging from surface-related features such as color and texture to shape-related features such as edge, curvature, and facial features — reflecting the MEI features we identified. Although Wang and colleagues did not study the columnar organization of V4, our combined results provide evidence for the hypothesis that the functional cortical columns we find are meticulously arranged into a cortical map.

### Similarity to tuning in deep networks

The MEIs for V4 neurons visually resemble MEIs of units in deep artificial neural networks trained on image recognition tasks. While there are differences between biological and artificial vision (reviewed in Sinz et al., 2019), deep networks trained on large-scale vision tasks are the closest human engineered system to biological visual system we know. Several previous works have found similarities between the primate visual system and deep network representations of visual stimuli (Güçlü & van Gerven, 2015; Khaligh-Razavi & Kriegeskorte, 2014; Yamins & DiCarlo, 2016; Yamins et al., 2014) and this similarity has recently also been shown in mice Bakhtiari et al. (2021); Nayebi et al. (2021). On a single neuron level, previous work has pointed out similarities between tuning in early vision and selectivities of single units in deep networks (Krizhevsky et al., 2012; Olah et al., 2020; Zeiler & Fergus, 2014) and a recent investigation has shown that single units in deep networks can exhibit similar object boundary tuning as V4 neurons (Pospisil et al., 2018). The striking similarity between our single cell MEIs of V4 neurons and single units in the InceptionV1 architecture (Olah et al., 2020) provide an even stronger case for similarities in tuning in primate early vision and deep networks. What’s more, by using this similarity we can derive predictions about the color selectivity of V4 neurons despite having shown only grayscale images in the experiment and during model training (Fig. 5b).

### Future directions

Our research highlights the power of applying deep learning to comprehensively characterize neuronal representations and to elucidate the relationships between brain structure and function. The concept of cortical columns offers an appealing framework for understanding cortical computation, as it decomposes complex processing into smaller, repeated building blocks. Building on prior evidence for columnar and clustered organization in sensory cortex, our findings support the hypothesis that groups of functionally similar neurons in the mid-level visual area V4 exhibit local clustering across cortical depth. Because neuronal connections within a cortical area tend to be locally dense Gilbert & Wiesel (1989), neighboring neurons within a column—both within and across layers— are much more likely to be interconnected. The purpose of a columnar architecture, therefore, is not to impose strict spatial boundaries between neurons, but rather to enhance synaptic opportunities among neurons that share similar response properties. Such local connectivity facilitates circuit computations that may underlie the emergence of new types of invariances or the construction of increasingly complex feature selectivity.

In this study, we focused on the most activating stimuli and on tuning similarities among neurons recorded along the same probe. However, recent work has extended this approach to include both the most and least activating stimuli, revealing that many visual cortical neurons encode information bidirectionally—as a contrast between two distinct feature types—rather than purely through excitation **?**. This bidirectional framework suggests that structured excitation and suppression jointly define functional micro-circuits, raising the intriguing possibility that such dual-feature coding may also manifest within columnar organizations in higher visual areas.

A comprehensive understanding of local cortical organization will therefore require characterizing full tuning functions—including invariances and the diversity of encoded features among nearby neurons. Future work combining large-scale functional recordings with synaptic-resolution connectomics Bock et al. (2011); Consortium et al. (2021); Ding et al. (2023a); Reid (2012) will be essential for clarifying how local circuits implement these computations.

## ACKNOWLEDGEMENTS

The authors thank the International Max Planck Research School for Intelligent Systems (IMPRS-IS) for supporting Konstantin Willeke and Arne Nix. The authors would also like to thank Jan Niklas Böhm, Philipp Berens, and Dmitry Kobak for their guidance with using their recently developed t-simCNE model and Greg Horwitz for feedback on the manuscript. The authors also thank Edgar Y. Walker, George Denfield, Christoph Blessing, Mohammad Bashiri, Konstantin-Klemens Lurz, Max Burg, Shanqian Ma, Robert Petrovic, Elena Offenberg and Paul Fahey for technical support and helpful discussions. The research was funded by the Carl-Zeiss-Stiftung (KW, FHS), the Cyber Valley Research Fund (AN, FHS). FHS is further supported by the German Federal Ministry of Education and Research (BMBF) via the Collaborative Research in Computational Neuroscience (CRCNS) (FKZ 01GQ2107), as well as the Collaborative Research Center (SFB 1233, Robust Vision) and the Cluster of Excellence “Machine Learning – New Perspectives for Science” (EXC 2064/1, project number 390727645). ASE received funding for this project from the European Research Council (ERC) under the European Union’s Horizon Europe research and innovation programme (Grant agreement No. 101041669) and Deutsche Forschungsgemeinschaft (DFG, German Research Foundation), project ID 432680300 (SFB 1456, project B05). The work was also supported by the Intelligence Advanced Research Projects Activity (IARPA) via Department of Interior/ Interior Business Center (DoI/IBC) contract numbers D16PC00003, D16PC00004, and D16PC0005. The U.S. Government is authorized to reproduce and distribute reprints for Governmental purposes notwithstanding any copyright annotation thereon. We also acknowledge support from the National Institute of Mental Health and National Institute of Neurological Disorders And Stroke under Award Number U19MH114830 and National Eye Institute award numbers R01 EY026927 and Core Grant for Vision Research T32-EY-002520-37 as well as the National Science Foundation Collaborative Research in Computational Neuroscience IIS-2113173. Disclaimer: The views and conclusions contained herein are those of the authors and should not be interpreted as necessarily representing the official policies or endorsements, either expressed or implied, of IARPA, DoI/IBC, or the U.S. Government.

## AUTHOR CONTRIBUTIONS

**KFW** Conceptualization, Methodology, Validation, Software, Formal Analysis, Investigation, Writing - Original Draft, Visualization **KR** Conceptualization, Methodology, Validation, Software, Formal Analysis, Investigation, Data acquisition, Data curation, Writing - Original Draft, Visualization; **KF** Conceptualization, Methodology, Investigation, Experimental and analysis design, Writing Original Draft, Writing - Review & Editing, Supervision; **AFN** Methodology, Software, Writing - Review & Editing; **SAC** Conceptualization, Methodology, Software, Investigation, Experimental and analysis design; **TS**,**CN**,**GR**,**SP** Data acquisition, Data curation, Methodology; **ASE** Conceptualization, Methodology, Investigation, Experimental and analysis design, Writing - Review & Editing, Supervision; **FHS** Conceptualization, Methodology, Investigation, Experimental and analysis design, Writing - Original Draft, Writing - Review & Editing, Supervision, Funding Acquisition, Project administration; **AST** Conceptualization, Methodology, Investigation, Experimental and analysis design, Writing - Original Draft, Writing - Review & Editing, Supervision, Funding Acquisition, Project administration;

## Materials and Methods

### Ethics statement

Electrophysiological data were gathered from a pair of healthy male rhesus macaque monkeys (*Macaca mulatta*), aged 17 and 19 years, and weighing 16.4 and 10.5 kg, respectively, at the time of the study. The research adhered to NIH guidelines and received approval from the Institutional Animal Care and Use Committee at Baylor College of Medicine (permit number: AN-4367). The monkeys were individually housed in a spacious room near the training facility, in the company of approximately ten other monkeys, allowing for abundant visual, olfactory, and auditory interactions. They were maintained on a 12- hour light/dark cycle.

The Center for Comparative Medicine at Baylor College of Medicine ensured that the monkeys received regular veterinary check-ups, balanced nutrition, and environmental enrichment. Surgical procedures involving the monkeys were performed using general anesthesia and adhering to standard aseptic techniques. Postoperative pain was managed with analgesics for seven days.

### Electrophysiological recordings

Non-chronic recordings were conducted using a 32-channel linear silicon probe (NeuroNexus V1x32-Edge-10mm-60-177), with surgical methods and recording protocols previously outlined (Denfield et al., 2018). In summary, custom titanium recording chambers and head posts were implanted under complete anesthesia and sterile conditions. Initially, the bone remained unaltered, and only before recordings were small trephinations (2 mm) made over lateral V4, with eccentricities spanning from 1.7 to 18.3 degrees of visual angle. Recordings took place after a week of each trephination. A Narishige Microdrive (MO-97) and guide tube were used to carefully lower the probes, penetrating the dura.

### Data acquisition and spike sorting

Electrophysiological data were continuously collected as a broadband signal (0.5Hz–16kHz), digitized at 24 bits. The spike sorting methods employed in this study resemble those used in (Cadena et al., 2019; Denfield et al., 2018), with the code accessible at https://github.com/aecker/moksm. The linear array of 32 channels was divided into 14 groups, each containing six neighboring channels (with a stride of two), which were treated as virtual electrodes for spike detection and sorting. Spikes were identified when channel signals exceeded a threshold equal to five times the standard deviation of the noise.

Following spike alignment, the first three principal components of each channel were extracted, resulting in an 18-dimensional feature space utilized for spike sorting. A Kalman filter mixture model was fitted to monitor waveform drift, which is common in non-chronic recordings (Calabrese & Paninski, 2011; Shan et al., 2017). Each cluster’s shape was modeled using a multivariate t-distribution (degrees of freedom = 5) with a ridge-regularized covariance matrix. The cluster count was determined based on a penalized average likelihood with a constant cost for each additional cluster (Ecker et al., 2014).

Lastly, a custom graphical user interface was used to manually confirm single-unit isolation by evaluating the units’ stability (based on drifts and cell health throughout the session), identifying a refractory period, and inspecting scatter plots of channel principal component pairs.

### Visual stimulation and eye tracking

Visual stimuli were generated by a specialized graphics workstation and presented on a 16:9 HD widescreen LCD monitor (23.8”) with a 100 Hz refresh rate and a resolution of 1920 × 1080 pixels, positioned at a 100 cm viewing distance (yielding approximately ∼63*px/*^*°*^). The monitor underwent gamma correction to ensure a linear luminance response profile. A custom-made, camera-based eye tracking system confirmed that monkeys kept their gaze within roughly ∼0.95*°* around a ∼0.15*°*-sized red fixation target. Offline evaluations revealed that the monkeys typically fixated with greater accuracy.

Upon maintaining fixation for 300 ms, a visual stimulus was displayed. If the monkeys sustained their gaze throughout the entire trial duration (1.8 s), they were rewarded with a drop of juice at the end of the trial.

### Receptive field mapping and stimulus placing

At the start of each session, we determined receptive fields in relation to a fixation target using a sparse random dot stimulus. A solitary dot, spanning 1*°* of the visual field, was displayed on a uniform gray background, with its location and polarity (black or white) randomly changing every 30 ms. Each fixation trial persisted for two seconds. Multi-unit receptive field profiles for each channel were obtained through reverse correlation. The population receptive field location was estimated by fitting a 2D Gaussian to the spike-triggered average across channels at the time lag that optimized the signal-to-noise ratio.

The natural image stimulus occupied the entire screen. The fixation spot was adjusted so that the mean of the population receptive field was as close to the screen’s center as feasible. Due to the recording sites’ location in both monkeys, this positioning involved placing the fixation spot near the screen’s upper border, shifted to the left.

### Natural image stimuli

We selected a collection of 24,075 images from 964 categories (∼25 images per category) from ImageNet (Deng et al., 2009), transformed them to grayscale, and cropped the central 420 × 420 pixels. For images that were smaller than 420 × 420, a central crop was taken and the resulting image was re-scaled to 420 × 420. Each image had an 8-bit intensity resolution (values ranging from 0 to 255). From this set, we randomly chose 75 images as our *test-set*. Out of the remaining 24,000 images, we designated 20% as *validation-set* at random, leaving 19,200 images in the *train-set*. Natural images were displayed during the standalone generation recordings of 1,244 units and during the generation phase of closed-loop recordings for 82 units. Specifically, *∼* 12k unique *train-set* images were displayed during the standalone generation recordings, and ∼7.5k unique *trainset* images were displayed during the generation phase of closed-loop recordings. Across sessions, train images were randomly sampled from the *train-set* such that the full set was exhausted before cycling back through the same images. The 75 *test-set* images were displayed in every recording session. Note that selecting images from ImageNet means that the pre-trained convolutional network (see below) has likely seen our natural stimuli (but not the neuronal responses) during training on the classification task.

During standalone generation recording sessions, ∼1000 successful trials (∼12k train images and 75 repeated test images) were recorded, whereas 600 successful trials (∼7.5k train images and 75 repeated test images) were recorded during the generation phase of closed-loop experiments. In both instances, each trial involved continuous fixation for 2.4 seconds, which includes 300 ms of a gray screen (intensity 128) at the beginning and end of the trial, as well as 15 consecutive images displayed for 120 ms each without any gaps. Trials contained either training/validation images that were presented only once or test images, which were repeated during the experiment.

Throughout the recording session, all trails were randomly interleaved, with test images being repeated 40-50 times during standalone generation recordings and 20 times during the generation phase of closed-loop recordings. Training and validation images were sampled without replacement, so each image was effectively displayed once or not at all. Images were upscaled using bicubic interpolation to match the screen width (1920 pixels) while maintaining their aspect ratio. The upper and lower 420-pixel bands were cropped out to cover the entire screen, effectively stimulating both classical and beyond classical receptive fields of V4 neurons. After sorting the neurons, spikes associated with each image presentation were counted within a 70-160 ms time window after stimulus onset.

## Image preprocessing for model training

Starting out from an original image size of 420 × 420 with a resolution of 14px*/*^*°*^ we cropped the upper and lower bands to fit the full screen for presentation for a resulting image size of 420 × 236. We then cropped the images so that only the bottom center 200×200 pixels remained. Then, we down-sampled the images to either 80 × 80 or 100 × 100 pixels (5.8px*/*^*°*^ or 7px*/*^*°*^), for the closed-loop model training and non closed-loop model training, respectively.

### Model architecture

Our neural predictive model of primate V4 consisted of two main parts: A pretrained *core* that computes nonlinear features of input images, and a *Gaussian readout* (Lurz et al., 2020) that maps these features to the neuronal responses of the single neurons.

As the core of our model, we used a ResNet50 (He et al., 2016) which was adversarially trained on ImageNet (Deng et al., 2009) to have robust visual representations (Salman et al., 2020), which yields improved transfer-learning performance (Engstrom et al., 2019a,b; Madry et al., 2017). Interestingly, it has been previously shown that robust features not only allow for better transfer-learning but they appear to be more similar to biological networks and also improve neural predictivity (Feather et al., 2022; Guo et al., 2022; Li et al., 2019; Safarani et al., 2021). Building on previous work (Cadena et al., 2022), we selected the first residual block of layer 3 of the ResNet, layer3.0, to read out from, and found that the adversarially robust training with *ϵ* = 0.1 yielded the highest predictive performance, compared to all other ResNet models and layers. The corresponding size of the output feature map at layer layer3.0 was 1024.

The input images **x** were forwarded through all layers up to a selected layer, to output a tensor of feature maps. Importantly, the parameters of the pretrained network were always kept fixed. We then applied batch-normalization (Ioffe & Szegedy, 2015). Lastly, we rectified the resulting tensor with a ReLU unit to obtain the final nonlinear feature space Φ(**x**) ∈ ℝ^*w×h×c*^ (**w**idth, **h**eight, **c**hannels) shared by all neurons.

To predict the response of a single neuron from the Φ(**x**) ∈ ℝ^*w×h×c*^ we use a *Gaussian readout* (Lurz et al., 2020). For each neuron *n*, this readout learns the coordinates (*x*^(*n*)^, *y*^(*n*)^) of the position of the receptive field on the output tensor and extracts a feature vector 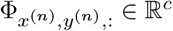 at this location from Φ. To this end, the Gaussian readout learns the parameters of a 2D Gaussian distribution 𝒩 (*µ*_*n*_, Σ_*n*_) and samples a location in feature space Φ(**x**) during each training step for every neuron *n*. Σ_*n*_ is initialized large enough to ensure that the entire visual field can be covered, and then decreases in size during training to have a more reliable estimate of the mean location *µ*_*n*_. At inference time (i.e. when evaluating our model), the readout is deterministic and uses the fixed position *µ*_*n*_. Although this framework allows for rotated and elongated Gaussian functions, we found that for our data, an isotropic formulation of the covariance – parametrized by a single scalar 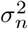 – was performing equally well as compared to a fully parametrized Gaussian. Taken together, total number of parameters per neuron of the readout were *c* + 4 (number of channels, bivariate mean, variance, and bias). This extracted feature vector 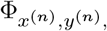 is then used in a linear-nonlinear model to predict the neuronal response. To this end, an affine function of the resulting feature vector at the chosen location was computed, followed by a rectifying nonlinearity *f*, chosen to be an ELU (Clevert et al., 2015) offset by one (ELU + 1) to make responses positive (Eq. 1). The weight vector **w**_*n*_ ∈ ℝ^*c*^ was *L*_1_ regularized during training.

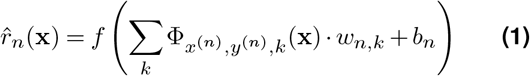

### Model training

We trained our model to minimize the summed Poisson loss across *N* neurons between observed spike counts *r* and the models’ predicted spike counts 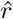 in addition to the *L*_1_ regularization of the readout parameters.

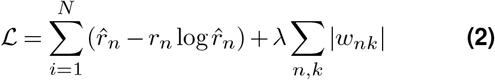

We trained the models either on the full dataset of n=100 recording sessions with n=1244 neurons and an image size of 100 by 100 pixels, or on an individual sessions during a closed loop recording with a reduced image size of 80 by 80 pixels in order to save time with model training and stimulus generation. For training on the full dataset, an epoch consisted of the cycling through the whole training set of 19200 images. However, because only *≈*9,000 - 13,000 images were shown in each session, if a particular image was not shown in a session, we simply zeroed out the gradients of all neurons in the sessions that did not contain the image in question. When training models on a single closed loop recording, we cycled through all *≈*9,000 training images that were shown in that session. In all cases, we used a batch size of 64 and after each batch, we updated the weights using the Adam optimizer (Kingma & Ba, 2014). The initial learning rate was 3 · 10^*−*4^ with a momentum of 0.1.

After each epoch, we computed the Poisson loss on the entire validation set. Similar to the training set, not all images were shown in all sessions, so that we again zeroed out the loss for the sessions in question. We then used early stopping to decide whether to decay the learning rate or stop the training altogether. We scaled the learning rate by a factor of 0.3 once the validation loss did not improve for five consecutive epochs. Before decaying the learning rate, we restored the weights to the best ones based on the poisson loss on the validation set. After four early stopping steps were completed, we stopped the training. On average, this resulted in *≈*50 training epochs, for a training time of 2 minutes for a closed loop session, and 15 minutes for the entire dataset on a NVIDIA 2080ti GPU.

## Ensemble models

Instead of using a single trained model, we used a model ensemble for all of our analyses and for MEI generation. To predict the neuronal responses to individual images, we trained readout weights for each member of an ensemble of five models initialized with different random seeds and used the average prediction across the ensemble for further analyses (Hansen & Salamon, 1990). We always trained ten individual models with a different random seed, which determined the model initialization as well as the drawing of training batches. Then, we selected the five models with the highest performance on the validation set to form a model ensemble. The inputs to the ensemble model were passed to each member, and the resulting predictions were averaged to obtain the final model prediction.

### Explainable variance

As a measure of response reliability, we estimated the fraction of the stimulus-driven variability as compared to the overall response variability. More specifically, we computed the ratio between each neurons’ total variance minus the variance of the observation noise, over the total variance (Eq. 3).To estimate the variance of the observation noise, we averaged the variance of responses across image repeats for all of the 75 repeated full-field natural image test stimuli 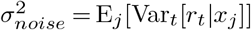 where *t* corresponds to the repeats and *x*_*j*_ represents a unique image:

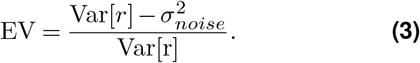

### Model performance measures

To measure the predictive performance of our models, we calculated the correlation to average (Cadena et al., 2022; Franke et al., 2022; Willeke et al., 2022) on the held out test set images. Given a neuron’s response *r*_*ij*_ to image *i* and repeat *j* and the model predictions *o*_*i*_, the correlation is computed between the predicted responses and the average neuronal response 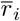 to the *i*^*th*^ test image (averaged across repeated presentations of the same stimulus):

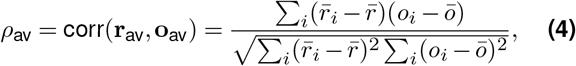

where 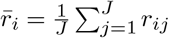 is the average response across *J* repeats, 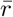 is the average response across all repeats and images, and 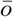 is the average prediction across all repeats and images.

### Generation of MEIs

We used the trained model to synthesize maximally exciting input images (MEIs) for each neuron using regularized gradient ascent (Bashivan et al., 2019; Franke et al., 2022; Walker et al., 2019). Starting out with a randomly initialized Gaussian white noise image given by **x** ∈ ℝ^*h×w*^, with height *h* and width *w*, we showed the image to the model and computed the gradients of a single target neuron w.r.t. the image. To avoid high frequency artifacts, after each iteration we applied Gaussian blur with a *σ* of 1 pixel to smoothen the image. Additionally, we constrained the entire image to have a fixed energy budget, which we implemented as a maximum L2 norm of the standardized image, calculated across all pixel intensities. We chose an L2 norm of 25 for all neurons such that the resulting MEIs had minimal and maximal values similar to those found in our training natural image distribution. If the MEI exceeded the allowed norm budget after any iteration, we divided the MEI by factor *f*_*norm*_ with *f*_*norm*_ = ∥MEI∥_2_*/b*. Additionally, enforced that the MEI could not contain values outside of the 8-bit pixel range by clipping the MEI outside of the bounds that correspond to 0 or 255 pixel-intensity. We used the stochastic gradient descent (SGD) optimized with learning rate of 10. We ran each optimization for 1000 iterations, without early stopping.

### MEIs with transparency masks

Furthermore, we employed a novel technique of synthesizing MEIs using a transparency channel based on the idea of Mordvintsev et al. (2018). Given an MEI optimized with a method as described in the section above, it is difficult to distinguish which of the MEI features are important, i.e. strongly activate the neuron, and which ones are not. Through our inbuilt L2 energy constraint, there only is finite contrast that the model is able to distribute across the image. However, this is still uninformative regarding which features of the MEI are most important. Thus, we adopted a transparency method as a differentiable parametrization (Mordvintsev et al., 2018), which jointly optimizes the MEI and a transparency mask, with the objective of making the MEI itself as transparent as possible (i.e. uninformative parts of the MEI), while still retaining the high neuronal activation of the resulting image. More specifically, we optimized an image **x** ∈ ℝ^*c×h×w*^, with channels *c*, height *h* and width *w*. We set the channels *c* = 2, treated the first channel as the MEI **x**_*mei*_, and the second channel as a transparent mask **x**_*α*_, which we optimized jointly as follows: (1) In each iteration, we drew a random image **x**_*bg*_ from the training set as a background. (2) We clip the second MEI channel, i.e. the transparent mask, between the values 0–1, with 1 meaning that the MEI is fully opaque, and 0 meaning the the MEI is not visible at all. (3) Then, we blend the background and the MEI according to the transparency mask to get the combined image **x**_*combined*_ with

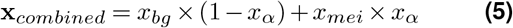

We then showed the combined image to the model to compute the neuronal response of the target neuron. With no additional constraints, the model did in fact learn to set all values of the transparent mask to 1, so that it is necessary to change the MEI objective function **L**_*old*_ from simply maximizing the neuronal response *r* to penalize alpha values of zero, as suggested by Mordvintsev et al. (2018):

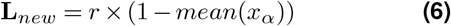

We used the same learning rate and optimizer as for our default gradient descent MEIs. We also applied the same L2 contrast constraint of 25, but only to the MEI image channel *x*_*mei*_. For our closed loop experiments, we ran the initial 16 sessions with the default gradient ascent MEIs, and the final 8 sessions with the transparency method, observing similar success for the *in vivo* verification. Because the transparency method seemed to generate MEIs that were less spread out and had features that were perceptually easier to identify, we used exclusively this method to generate the MEIs for the full dataset. For all the 889 out of 1244 neurons that passed our threshold of correlation to average larger than 0.3, we generated 50 MEIs per neuron for a total of 44,450 MEIs. Each MEI took on average 2 minutes on a NVIDIA 2080ti GPU, resulting in a total compute time of *≈*60 GPU weeks.

### Centering of MEIs

In this work, we do not further analyze the transparent MEI masks. However, we use the transparent masks to center the MEIs for the similarity quantification. After MEI optimization, the MEI is located at the approximate RF location of each neuron. To center the MEIs, we find the center of mass of the transparent mask, and move the MEI such that the center of mass is at the center. All MEIs shown throughout this manuscript are shown at the center location for ease of comparison.

### Psychophysics

In our psychophysics experiment,the objective was to assess the perceptual similarity between MEIs originating from either a single recording session as compared to a random selection of neurons from different recording sessions. For this purpose, we devised a two-alternative forced choice task in which we showed two sets of MEIs, arranged in 3x3 grids as the basis for the similarity evaluation task. Participants were instructed to simply report which of the MEIs in the two grids looked more similar to each other. No other information apart from the displayed MEIs was provided.

To obtain the images for the task, we randomly chose 50 out of 52 sessions that had a minimum of 9 well-predicted neurons, using the threshold of correlation to average larger than 0.3. From these selected sessions, the top 9 well-predicted neurons were identified, resulting in a total pool of 450 neurons. For each session, participants were presented with images from the top 9 predicted neurons and a randomly selected set of 9 additional neurons from the pool, sampled without replacement. Specifically, for each neuron, we presented five MEIs, generated by starting from a different initial noise image during optimization, as images and compiled in a GIF. Participants were shown the two example sessions which were not included in the study, together with the correct solution. The subjects had unlimited time to make their choice.

Subject recruitment involved lab members that were familiar with the concept of an MEI from all research groups contributing to this article. In total, we recorded the responses of N=25 observers.

### Closed-loop procedure

Our closed loop experiments were composed of a generation and verification session. In the generation session, we recorded the electrophysiological responses of V4 neurons to full-field gray-scale natural images for 600 trials, corresponding to 9,000 unique images. After showing of all of these trials, we stopped and promptly restarted the neurophysiological acquisition system to continue recording neuronal activity while we processed the data from the generation session. During this analysis period, we isolated single units and used these units to train several models to predict neuron responses to natural images. We produced an ensemble of the best five models, used this ensemble to select the six highest predicted units and generate ten MEIs per unit. We computed the mask of each MEI by calculating the z-score of each pixel value and setting a threshold (*z* = 0.35) to isolate the area within the mask. We then created a convex hull to close any holes and smoothed the edges with a Gaussian filter (*σ* = 2). We used this masking procedure for our natural image control selection procedure.

Specifically, we screened a distinct set of 5,000 natural images, different from our training, validation, and test set images. This set was selected randomly from a database of 100k images, and we screened this same image set for all neurons across all closed-loop experiments. Each image was masked with the ten MEI masks of each unit and re-normalized to the contrast of the MEIs, resulting in 50,000 total screened images for each unit. We selected the seven highest activating masked natural images of each unit to use as controls for the verification recording.

### Closed-loop stimulation paradigm

During the verification session of the closed-loop experiment, we displayed one MEI and seven controls for each of the six units. These stimuli were centered on the RF and repeated 10 times throughout the experiment. We also showed the same full-field test set natural images 20 times each, to enable functional response matching between the generation and verification units. Each trial contained 15 images, composed of either (i) randomly interleaved generated stimuli (i.e. MEIs and masked controls) or (ii) full-field test set natural images, as described previously. Trials were randomly interleaved. In total, 100 trials of full-field images and 180 trials of generated stimuli were shown.

### Closed-loop unit matching

To assess the stability of our closed-loop neurons, we developed a procedure for spike waveform matching single units between the generation and verification session recordings. To do this, we performed a *post hoc* spike sorting of the full recording session, which was produced by stitching together the raw data files of the generation and verification recordings. The verification recording includes the analysis period immediately after the generation recording, during which we isolate single units and generate their MEIs. This continuity of recording allows us to more easily track single units throughout the experiment. Furthermore, spike sorting the full session allows us to use a drift correction of the spike sorting model based on a kalman-filter across the entire experiment, which aided the accuracy of our single unit tracking.

We then executed a two-step matching procedure using the spike sorting results of both the generation and full sessions. To determine potentially matched units, we first took the spikes of the generation session units and assigned them to full session units. We then calculated the proportion of spikes that were assigned exclusively to one unit in both the generation and verification sessions. If a unit in the generation session had at least 95% of its spikes assigned to only one single unit in the full session, this was considered a possible match. To confirm true matches, we assigned the full session spikes to the generation session units. If a unit in the full session had at least 95% of its spikes assigned to the potentially matched single unit in the generation session, these units were verified to be a match, and thus certified as stable. We additionally assessed neuron stability by computing the functional consistency of each unit. To compute this measure, we used the test set of 75 full-field natural images that were shown in both generation and verification session and calculated the Pearson correlation to the repeat-averaged responses in both sessions. We set a minimum functional consistency threshold of 0.5. Taken together, we were able to waveform match 82 neurons from 24 closed-loop sessions, with 27 of these failing to meet the functional consistency criterion, resulting in n=55 neurons for the analysis of the closed-loop paradigm.

### MEI similarity quantification using contrastive learning

We used a recently proposed method of contrastive learning (Böhm et al., 2023) to quantify the perceptual similarity of our neurons’ MEIs. The method is based on SimCLR (Chen et al., 2020), a method of self-supervised contrastive learning of visual representations in which images are embedded in a high-dimensional space. In this space, augmentations of the same image are learned to have small distances while simultaneously training the model to increase the distance to all other images. Böhm et al. (2023) adapted this procedure for a two-dimensional space by combining the ideas of contrastive learning with 2-d neighbor embeddings as used in t-SNE (van der Maaten & Hinton, 2008). Their new method, called tSimCNE, achieves this by changing the similarity function between image embeddings from cosine-similarity used in SimCLR, which would constrain the embeddings to the unit circle, to Euclidean distance *d*_*ij*_ = ∥**z**_*i*_ − **z**_*j*_ ∥ between the embeddings **z**_*i*_ and **z**_*j*_ of two augmentations *i* and *j* of the same image. We use this method to train a model end-to-end to embed our MEIs using the loss function as proposed by Böhm et al. (2023):

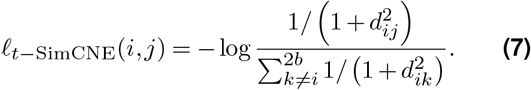

In our adaptation, *i* and *j* correspond to two data augmentations of the same MEI, and *z* denotes the 2-d embedding, which is the model output. For a given batch size *b*, the resulting batch size is 2*b* because each MEI is augmented twice. As pointed out by the authors of SimCLR (Chen et al., 2020), this approach requires careful selection of data augmentations, as well as choosing the largest batch size possible.

Taken together, our training procedure consists of these steps: We first select all 889 neurons above the model performance threshold of correlation to average larger than 0.3, as described earlier. For each neuron, from the 50 MEIs that we optimized per neuron, we select only the MEIs that elicited at least 90% of the predicted firing rate of the highest-acivating MEI, which resulted in anywhere from 3 to 42 MEIs per neuron, with a total of n=19,688 MEIs used in this and subsequent analyses. This selection was important as there can be failures during optimizing MEIs, resulting in noisy images which the contrastive learning algorithm was very sensitive to. Next, from all the MEIs that we selected, we assembled a batch of size n=889, by randomly drawing one MEI for each neuron.

Subsequently, we preprocessed the MEIs and applied the data augmentations in the following order: (1) centering the MEIs as described above, (2) center cropping to resize the MEIs from 100×100 to 75x75 pixels, (3) random rotation between 0 and 30 degrees, (4) random rescaling between 32×32 and 75×75 pixels, (5) random cropping from the resulting image to a final size of 32x32 pixels. Steps (2) to (4) were applied twice to each MEI to obtain two augmentations per MEI. The resulting training batch thus consisted of a randomly drawn MEI per neuron, augmented twice, for a total batch size of 889*2=1,778 images, with the model’s objective being to minimize the distance between each pair, and to maximize the distance from all-to-all images which are not pairs. It is important to point out that with this training scheme, the trained model was given no information about recording sessions or which MEIs belong to which neuron. Once the model was fully trained, we obtained the full 2-d embedding of all MEIs from all neurons by applying transformations (1) and (2), i.e. centering and center cropping the MEIs, as well as rescaling the MEIs to 32x32 pixels, and subsequently showing the resulting images to the model. Finally, We obtained a 2-d location of each neuron by taking the mean location over the embedding locations of all MEIs per neuron.

### Contrastive learning: model architecture and training

We closely followed the t-simCNE authors (Böhm et al., 2023) in our choice of model architecture and training paradigm. As a model backbone, we employed a randomly initialized ResNet18 (He et al., 2016) with an output size of 512 and a reduced kernel size of the first convolutional layer from 7 *×* 7 to 3 *×* 3. Following the authors, we also added one hidden ReLU layer with n=1,024 units followed by a linear output layer of n=128 units, which was reduced to n=2 during the final stage of training. The training consisted of three stages: First, the model was trained for 3,000 epochs with the output layer size of 128. In the second stage, the output layer was disregarded and replaced with a linear output layer of n=2, followed by a training of only this layer for 200 epochs while the rest of the model was frozen. Lastly, in the third stage, the whole model was fine-tuned for another 1,000 epochs with a reduction of the learning rate by a factor of 1,000. In each of the three stages, we used the initial learning rate of 0.03·*b/*256 *≈* 0.1 with b=889, preceded by linear warm-up for ten epochs, followed by cosine annealing (Loshchilov & Hutter, 2016) with a final learning rate of 0. In each epoch, we accumulated the loss from 10 batches and optimized the model using SGD with a momentum of 0.9.

### Contrastive learning: within-neuron MEI distances

As mentioned above, it is crucial that our contrastive learning training scheme did not provide the model with any information about either the recording sessions or the neurons’ identity of each MEI. Therefore, an important sanity-check for the trained model is to analyze the 2-d embedding distances of all MEIs that we optimized for a single neuron. We performed this analysis by simply calculating all pairwise distances and taking the average of all MEIs per neuron.

### Contrastive learning: Session distances

We first averaged all pairwise distances across neurons recorded within one session to obtain a within-session distance in the 2-d embedding space. We then compared this within-session distance to a shuffled control. For this control, we used bootstrapping to sample at random the same number of neurons as in a given session, with the constraint that they cannot be from the session in question and that all randomly drawn neurons originate from different sessions. We then computed the mean pairwise distance for these shuffled neurons, and repeated this process 25,000 times for each session to get the null distribution. We then calculated the percentile of the true within-session distance against each sessions’ null distribution to obtain a p-value.

### Contrastive learning: HDBSCAN cluster cutting

After we obtained the 2-d embedding space of MEIs, we clustered this entire two-dimensional space using hierarchical density-based spatial clustering of applications with noise (HDBSCAN McInnes et al., 2017), the same clustering that was employed by the t-simCNE authors (Böhm et al., 2023). We searched over the parameter grid minimum cluster size ∈ {100, 125,…, 400} × and minimum samples, which indicates the minimum number of samples to be considered non-noise, 5, ∈ {15,…, 145} with the constraint min-samples ≤ min-cluster-size. The resulting clusterings will create one clustered group labelled -1, for MEIs that were unable to be clustered. We selected the parameters min-cluster-size = 200 and min-samples = 10 which resulted in the lowest number of unassigned MEIs. Using this approach, the MEIs of each neuron get assigned a label, with the possibility that not all MEIs of single neuron were assigned to the same cluster. We assigned a cluster ID to a neuron if more than half of its MEIs had the same cluster ID. Furthermore, we excluded neurons for which the more than half of the MEIs were in the unassigned category -1 of the HDBSCAN algorithm. Based on these two criteria, 828 out of 889 neurons were given a valid clustering ID.

### Representational similarity of MEIs

As an independent way to verify the similarity of MEIs, we computed their representational similarity. For this purpose, we used the centered MEIs, and computed the responses of all 889 well-predicted neurons from our neural predictive model to each MEI. We controlled for the different RF positions of each *in silico* neuron by artificially centering the *in silico* neuron’s RF to the center by setting the readout location of the Gaussian readout to the center of the core output. We used this approach for each model ensemble member individually and again took the model ensemble average as the predicted neuronal activity. Consequently, for each MEI, we obtained a response vector of length 889, which we then used for pairwise comparisons using cosine similarity. We then compared the average cosine similarity either within clusters or within recording sessions to the average similarity across sessions/clusters. To obtain a p-value for the similarity within vs. across clusters, we computed the similarity matrix 100 times. In each of the 100 runs, per neuron we drew at random one of the neurons MEIs and computed the population response vector for that MEI. We thus ended up with 100 similarity matrices, for which the pairwise comparisons were thus computed based on different MEIs. For each cluster, we then computed the average similarity across the 100 runs, and compared this similarity. We then took (1) the average cosine similarity within each cluster across the 100 runs, and (2) for each of the 100 runs, for each cluster we average the similarity to all other clusters. We report the p-value as percentile of the within-session similarity to the distribution of across-session similarity.

## Data and code availability

The analysis code will be publicly available in an online repository latest upon journal publication. Our coding framework uses Pytorch (Paszke et al., 2019), Numpy (Harris et al., 2020), scikit-image (Van der Walt et al., 2014), matplotlib (Hunter, 2007), seaborn (Waskom et al., 2017), DataJoint (Yatsenko et al., 2015), Jupyter (Kluyver et al., 2016), and Docker (Merkel, 2014). We also used the following custom libraries and code: neuralpredictors (https://github.com/sinzlab/neuralpredictors) for torch-based custom functions for model implementation, nnfabrik (https://github.com/sinzlab/nnfabrik) for automatic model training pipelines using DataJoint, nnvision (https://github.com/sinzlab/nnvision) for specific model definitions, analysis, and figures.

## Supplementary Information

Supplemental Fig. 1 - Waveform and functional matching of single units across recordings

Supplemental Fig. 2 - Diverse model-derived stimuli of individual monkey V4 neurons

Supplemental Fig. 3 - Similarity of optimal stimuli in neuronal response space

Supplemental Fig. 4 - Overview of optimal stimuli of V4 response modes

**Supplemental Fig. 1.**
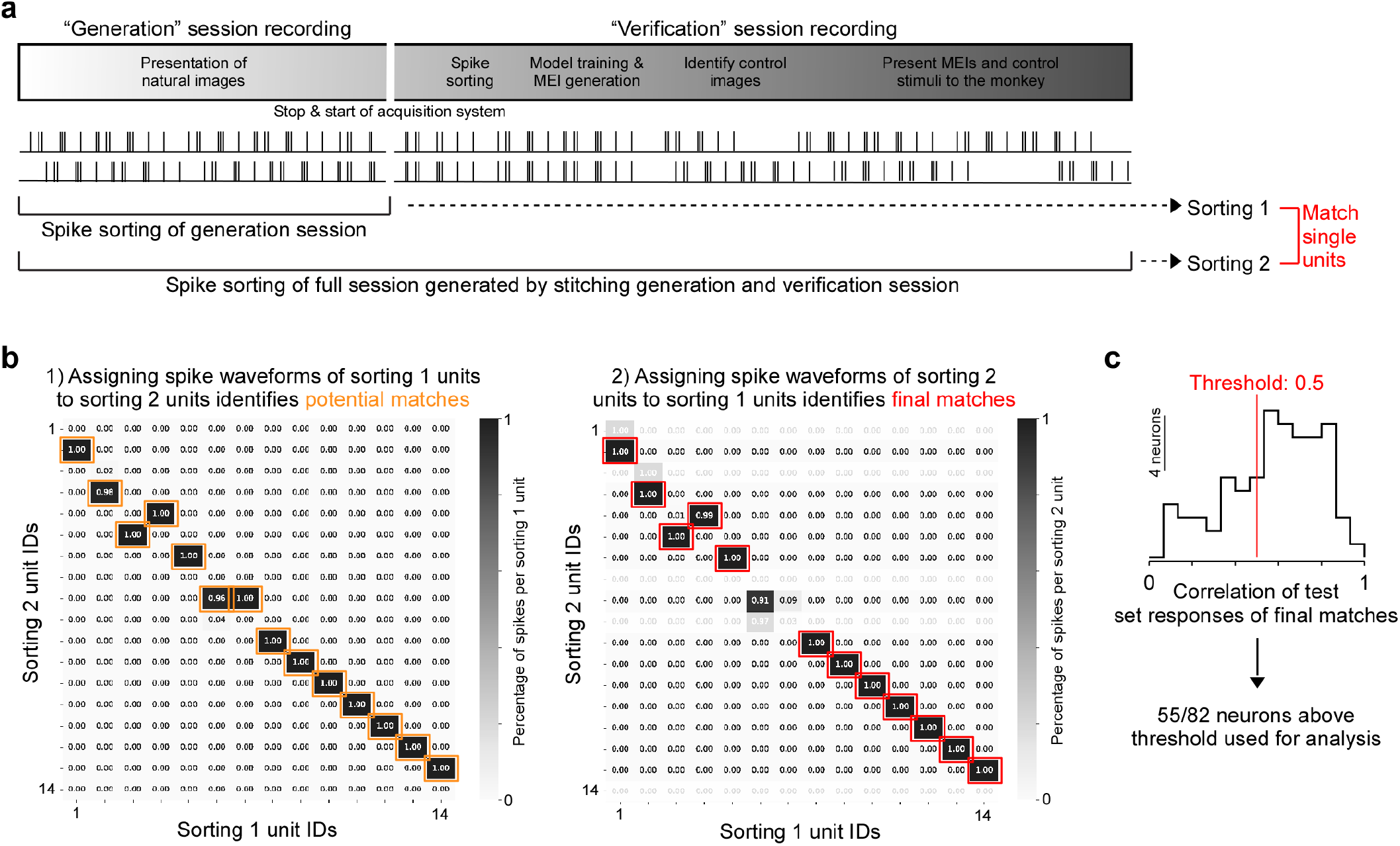
Spike waveform and functional matching of single units across recordings. **a**, Schematic illustrating spike sorting of the closed-loop experimental paradigm. The “generation session” is spike sorted directly after the recording, resulting in “Sorting 1”. This data is then used for model training and optimization of MEIs, which are presented back to the animal during the “verification session”. The verification session recording starts immediately after the generation session recording ends, to ensure a continuous monitoring of the recorded units over time. After the experiment, the generation and verification session recordings are concatenated (“full session”) and spike sorted, resulting in “Sorting 2”. **b**, Unit matching based on spike waveforms across Sorting 1 (generation session) and Sorting 2 (full session) for an example session. The left plot shows the percentage of spikes of the Sorting 1 units assigned to the units of Sorting 2. Units were assigned by passing the principal components of each spike, extracted using the Sorting 1 Gaussian Mixture model (GMM), to the Sorting 2 model. For a potential match (orange), at least 95% of the spikes of a single unit of Sorting 1 had to be assigned to an individual unit of Sorting 2. The right plot shows the percentage of spikes of Sorting 2 units assigned to the units of Sorting 1. For a final match (red), at least 95% of the spikes of a single Sorting 2 unit had to be assigned to the potential match of Sorting 1. **c**, Distribution of correlations of mean test set responses for all final matches. We only included matched units into the analysis, if their functional correlation was at least 0.5.

**Supplemental Fig. 2.**
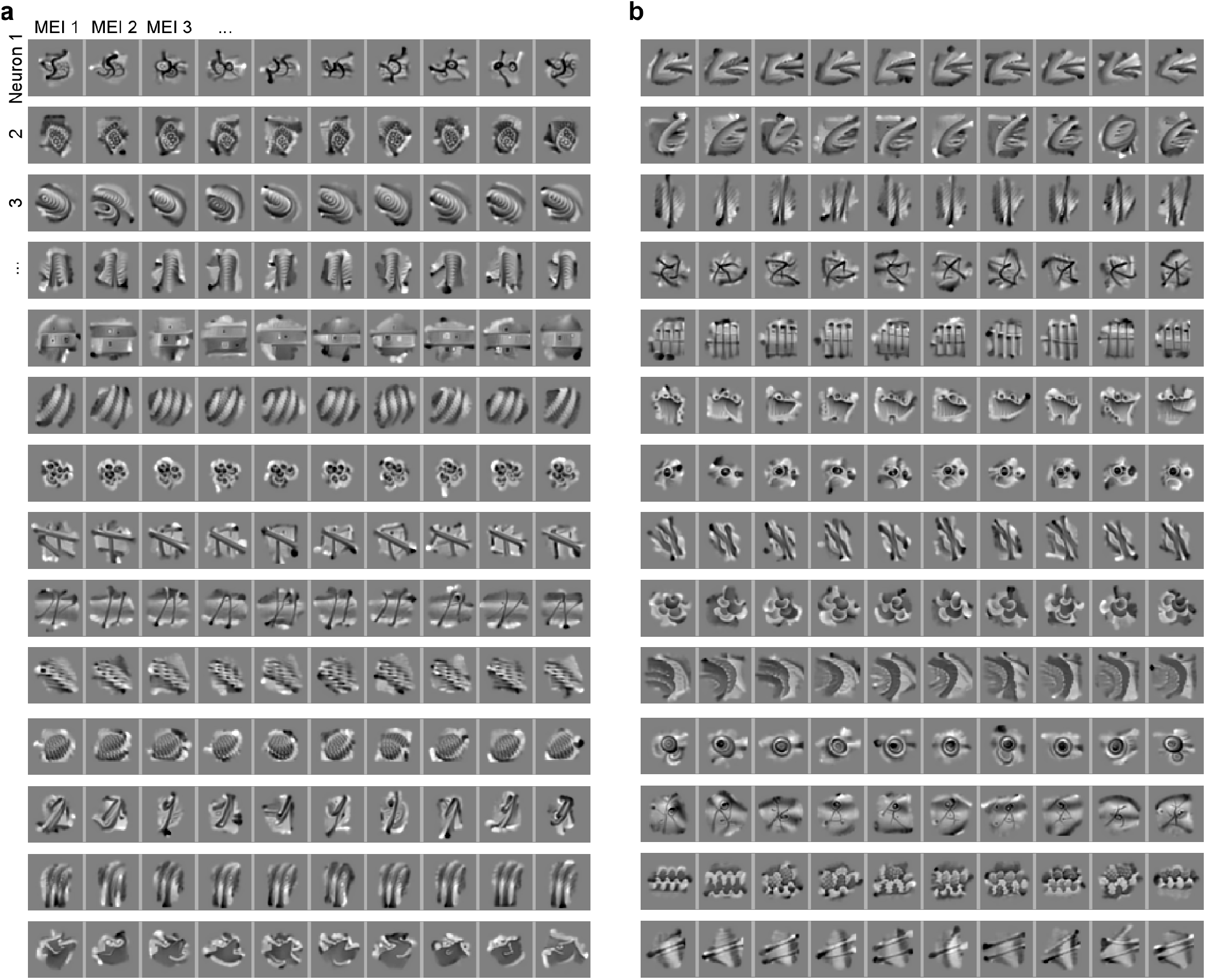
Diverse model-derived stimuli of individual monkey V4 neurons. **a**, For a set of 14 example neurons, we show 10 MEIs per neuron, generated from different starting points (random seeds) during the MEI optimization. The different MEIs exhibit the same visual feature but somewhat differ with respect to orientation, scale and position. **b**, Same as (a), for another set of 14 neurons.

**Supplemental Fig. 3.**
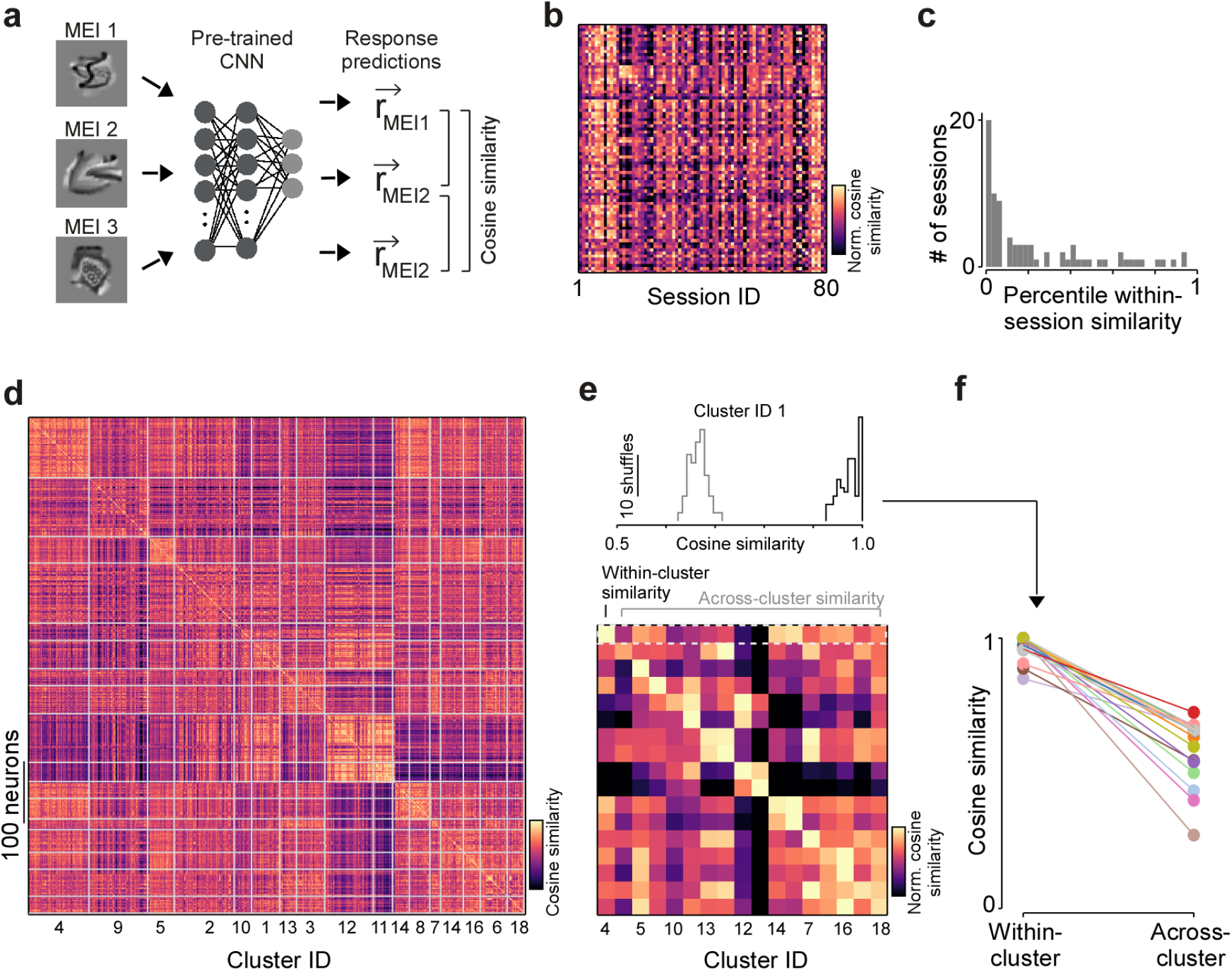
Similarity of optimal stimuli in neuronal response space. **a**, MEIs of three example neurons. Right shows a schematic illustrating how we compared the similarity of MEIs using representational similarity. In brief, each MEI was presented to the trained CNN model that was used to produce the MEIs to obtain a response vector. The response vectors were then compared using cosine similarity. **b**, Mean cosine similarity of MEIs within a single recording session (diagonal) and across recording sessions for n=88 sessions, peak-normalized for each row. **c**, Distribution of percentiles of within-session similarity. For example, a percentile of 0.05 means that the MEI similarity within the session was larger than the MEI similarity to 95% of the other sessions. For n=30/55 sessions, the percentile was <0.05. **d**, Cosine similarity of MEIs of n=889 neurons, sorted based on cluster assignment (cf. Fig. 5). **e**, Mean cosine similarity of MEIs within a cluster (diagonal) and across clusters, peak normalized per row. The matrix depicts the mean across n=100 similarity matrices, each generated based on a different random selection of MEIs per neuron. Top shows distribution of cosine similarity within an example cluster (black) and the mean similarity to all other clusters (gray). **f**, Mean within-cluster and across-cluster similarity for all clusters.

**Supplemental Fig. 4.**
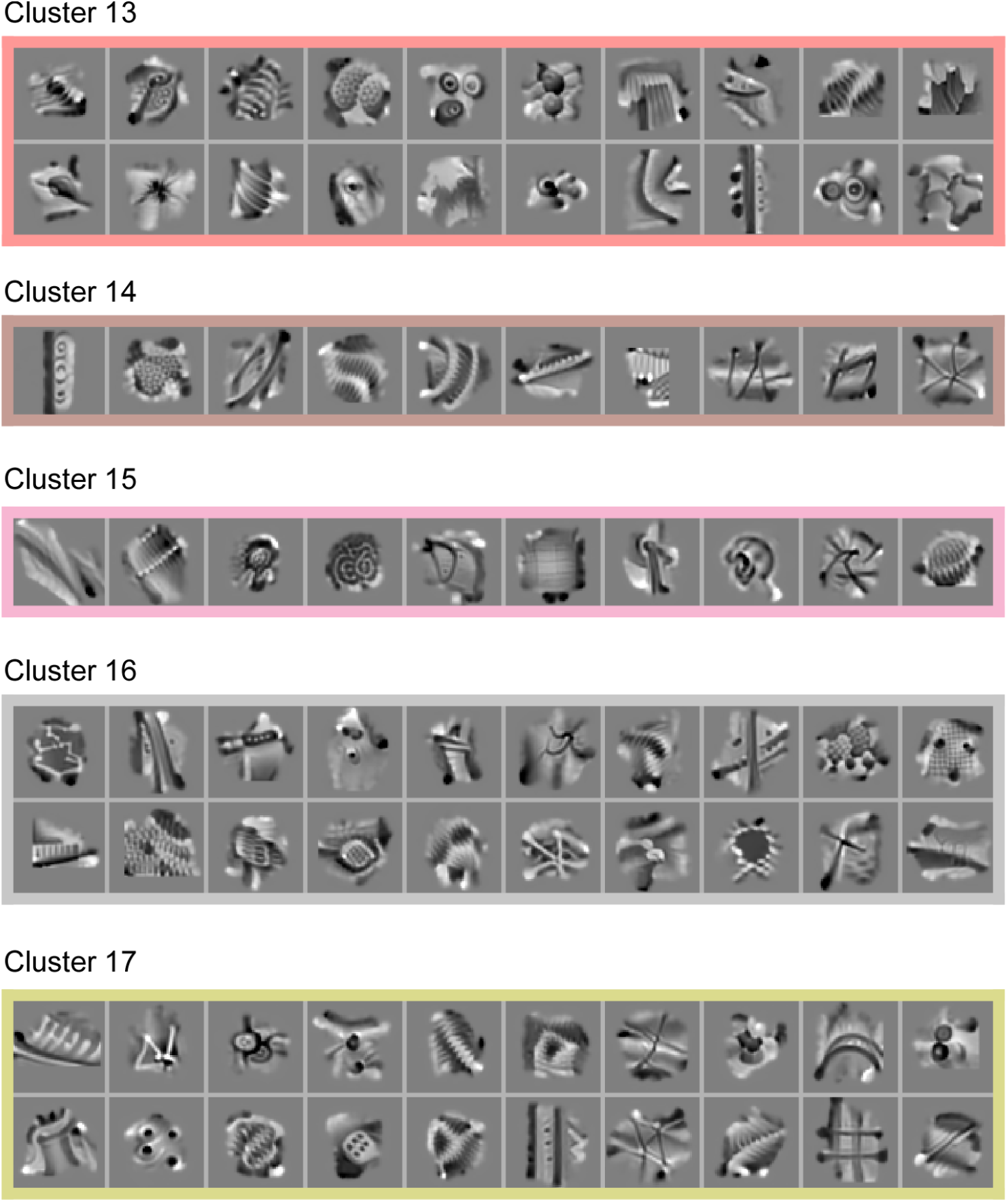
Overview of optimal stimuli of V4 response modes. **a**, MEIs of example neurons for five response modes not shown in Fig. 5.

